# Convergent Gene Expression Patterns During Compatible Interactions Between Two *Pseudomonas syringae* Pathovars and a Common Host (*Nicotiana benthamiana*)

**DOI:** 10.1101/2022.06.26.497614

**Authors:** Morgan E. Carter, Amy Smith, David A. Baltrus, Brian H. Kvitko

## Abstract

Summary/Abstract

*Pseudomonas syringae* is a diverse phytopathogenic species complex, and includes strains that can cause disease across a wide variety of plant species. Much previous research into the molecular basis of immunity and infection has focused on pathogen and plant responses in a handful of model strains and hosts, and with a tacit assumption that early steps in infection and host resistance are generalizable to the species complex and across plant hosts as a whole. Here, we provide a test of this assumption by measuring the dual pathogen and host transcriptomes of two distinct pathogenic lineages of *P. syringae* during compatible infection of a shared model host (*Nicotiana benthamiana*). Our results demonstrate that, with a handful of exceptions, host plants largely respond in a similar way to both pathogenic lineages and both bacterial pathogens possess highly similar transcriptional responses at 5 hours post inoculation. However, we also highlight that subsets of genes with differential expression patterns in both bacteria and host which likely represent strain-specific responses.

## Introduction

Knowledge of virulence mechanisms and strategies of phytopathogens has grown dramatically over the past few decades, largely due to a focus on gaining a deep understanding of infection in a handful of well-vetted and tractable strains and host plants. As one steps out to lesser studied strains and hosts, it is often assumed that relatively closely related bacterial phytopathogens use similar virulence pathways and strategies to overcome host defenses to cause disease. While such assumptions are rightly borne out of a desire for experimental efficiency and often hold up to scrutiny, they nonetheless remain assumptions until proven otherwise even if pathogens differ in timing and symptom development across hosts. With this idea in mind, we sought to categorize similarities and differences in responses of a common host plant to closely related bacterial phytopathogens during the early stages of infections under compatible interactions.

*Pseudomonas syringae* sensu lato consists of a collection of bacterial phytopathogens spanning a variety of formal species names and which is composed of upwards of 50 different pathovars (Susan S. Hirano and Upper 2000; Baltrus, McCann, and Guttman 2017). Pathovar designation for *P. syringae* sensu lato strains is typically designated based on phenotypic information such as host of isolation and more recently informed by genotypic, genomic, and phylogenetic characteristics (Baltrus, McCann, and Guttman 2017; Baltrus 2016). Much research effort has been spent to develop multiple *P. syringae* strains as model systems for virulence in a variety of host plants and these studies have been foundational for understanding bacterial virulence strategies as well host resistance to these pathogens (Lindeberg et al. 2006; Block and Alfano 2011; Xin, Kvitko, and He 2018). However, despite the incredible accumulation of data about biology and genomics of specific *P. syringae* strains, much of our understanding of virulence patterns remains dependent on assumptions of similarity of virulence strategies across various pathovars within the *P. syringae* sensu lato complex or is the product of indirect comparisons of infection by different pathovars on diverse and non-overlapping suites of hosts. We therefore sought to compare and contrast host plant responses and the infection strategies across two *P. syringae* strains that can each infect and cause disease (albeit with different symptom trajectories) on *Nicotiana benthamiana*.

We focused on two closely related strains for this study: *P. syringae* pv. *syringae* B728a (hereafter *Psy*) and *P. amygdali* pv. *tabaci* 11528 (hereafter *Pta). Psy* is a member of phylogroup 2 and was originally isolated as the causative agent of brown spot disease in common bean plants (*Phaseolus vulgaris*), and is well studied for its capabilities of survival as a plant epiphyte and and as a pathogen of numerous hosts (Feil et al. 2005; Helmann, Deutschbauer, and Lindow 2019; Baltrus, McCann, and Guttman 2017). *Psy* has also been widely to study bacterial infection of *N. benthamiana* under laboratory conditions(Vinatzer et al. 2006; Mohr et al. 2008; Misas-Villamil, Kolodziejek, and van der Hoorn 2011). *Pta* is a member of phylogroup 3 and was originally isolated as the causative agent of wildfire disease in tobacco plants (*Nicotiana tabacum)* but has also been reported to cause similar disease across bean hosts (Sun et al. 2021; Baltrus, McCann, and Guttman 2017). Previous genomic comparisons of these strains in the context of *P. syringae* diversity have suggested that these strains may differ in virulence strategies (Baltrus et al. 2011; Hockett et al. 2014); while a type III secretion system is critical for infection of hosts by both strains (S. S. Hirano et al. 1999) the effector repertoire of phylogroup 2 strains including *Psy* (16 effectors) appears reduced compared to that of many other analyzed strains with this reduction strongly correlated with acquisition of a suite of phytotoxins (syringomycin, syringopeptin, and syringolin) (Hockett et al. 2014; Baltrus et al. 2011). *Pta* shares 8 effectors with *Psy* and maintains a slightly larger repertoire than *Psy* (18 unique effectors and 21 total, reannotated herein, Supplemental File S1), but also contains a different suite of phytotoxins (tabtoxin and phevamine). Despite the potential for differences in virulence strategies, to date there have been no direct comparisons of host responses to these strains during compatible infection.

We investigated the two compatible disease interactions between *N. benthamiana* and *Psy* or *Pta* at the early time point of five hours post inoculation to understand how a plant host responds to two related bacteria with different infection strategies. Our results suggest that both pathogenic lineages of *P. syringae* display largely conserved patterns of transcription during the early stages of infection. We also highlight that, while there was clear overlap in the differentially expressed genes in *Nicotiana* in response to both pathogens, there were also clear transcriptome responses related to hormones and chloroplasts that were unique to infection by a specific strain.

## Results

### A complete genome sequence for P. syringae pv. tabaci ATCC11528 and reannotation of virulence factors

We and others have previously reported draft genome assemblies for *P. syringae* pv. tabaci strain ATCC11528 (Baltrus et al. 2011; Studholme et al. 2009), and here we report a complete genome assembled using a hybrid strategy that combined Illumina and Nanopore reads. The genome contains one circular chromosome that is 6,133,558 bp (Genbank accession CP042804.1) and one 68,162 circular plasmid (Genbank accession CP042805.1) which we name pTab1. The genome is predicted to encode a total of 5,488 proteins, 66 tRNA loci, 5 complete rRNA operons and 4 other non-coding RNA. We have used this genome sequence to reanalyze and revise annotations of type III effector proteins and their regulatory *hrp*-boxes. These annotations can be found in Supplemental File S1.

We have used the complete genome sequence for strain *Pta* to update annotations and positions of known virulence factors. We previously noted *hopR* and *hopAB* as possibly truncated and/or incomplete (Baltrus et al. 2011), and this was shown to be an error when different versions of this genome were reported. We note here that both *hopR* and *hopAB* appear to be full length and non-truncated in this complete genome assembly. We have confirmed that *hopAA1*, *hopAH1*, and *hopAI1* all appear to be truncated via nonsense mutations within this complete genome. Lastly, we find that this genome contains four identical copies of the effector *hopW*, denoted *hopW*1-1 through *hopW*1-4 in different positions throughout the chromosome and plasmid. Indeed, *hopW*1-4 is the only type III effector found on plasmid pTab1. In total, this strain is predicted to encode 18 full length type III effectors. We also note positions of the two phytotoxins known or predicted to be produced by this strain. Tabtoxin production is determined by two separate operons found in proximity to each other on the genome. However, surprisingly, we find that there are two regions of the chromosome that contain genes implicated in producing phevamine and that these two regions are identical in nucleotide sequence. We have labeled the first phevamine operon 1, and refer to these three genes as *hsv*1-1, *hsv*2-1, and *hsv*3-1. Genes within the second phevamine operon are labeled as *hsv*1-2, *hsv*2-2, and *hsv*3-2.

### RNA-seq analysis showed differential plant-gene expression patterns in response to both pathogens

We have included a schematic of our RNA sampling scheme as Supplemental Figure S1. This figure reflects the procedure carried out for one biological replicate and three biological replicate infections were carried out for each strain as part of this RNAseq experiment. RNA was collected and sequenced from two sets of *N. benthamiana* plants before inoculation and five hours post inoculation (hpi) with either *Pta* or *Psy*. Transcriptomes were compared for plants before and after inoculation and the threshold for differential expression was set at an adjusted p-value (padj) of 0.01 and an absolute value of the Log2 Fold Change (L2FC) above 2 (**Figure 1A**, **Supplemental Figure 3, Supplemental Table 1**). A set of 2245 genes (**Figure 1B**) are differentially expressed (DE) in plants in response to both pathogens by plants when comparing 0 hpi to 5 hpi. Although the 5 hpi time point represents genes with transcriptional responses both to infection by the *Pseudomonas* pathogens as well as genes that might have changed expression over the course of the 5 hour infection but which are unrelated to pathogen responses (such as the wound response), we still observed informative trends. Gene ontology (GO) enrichment analysis yielded 138 enriched GO terms in the shared DE genes (**Supplemental Figure 3**), with response to wounding at the bottom of the biological processes list. Many of the top enriched GO terms across categories are related to photosynthesis and the response to light, (i.e. GO:0016168 chlorophyll binding GO: 0015979, photosynthesis, GO: 0009579 thylakoid, etc.) with the associated genes being largely downregulated (**Figure 2**).

**Figure 1.**
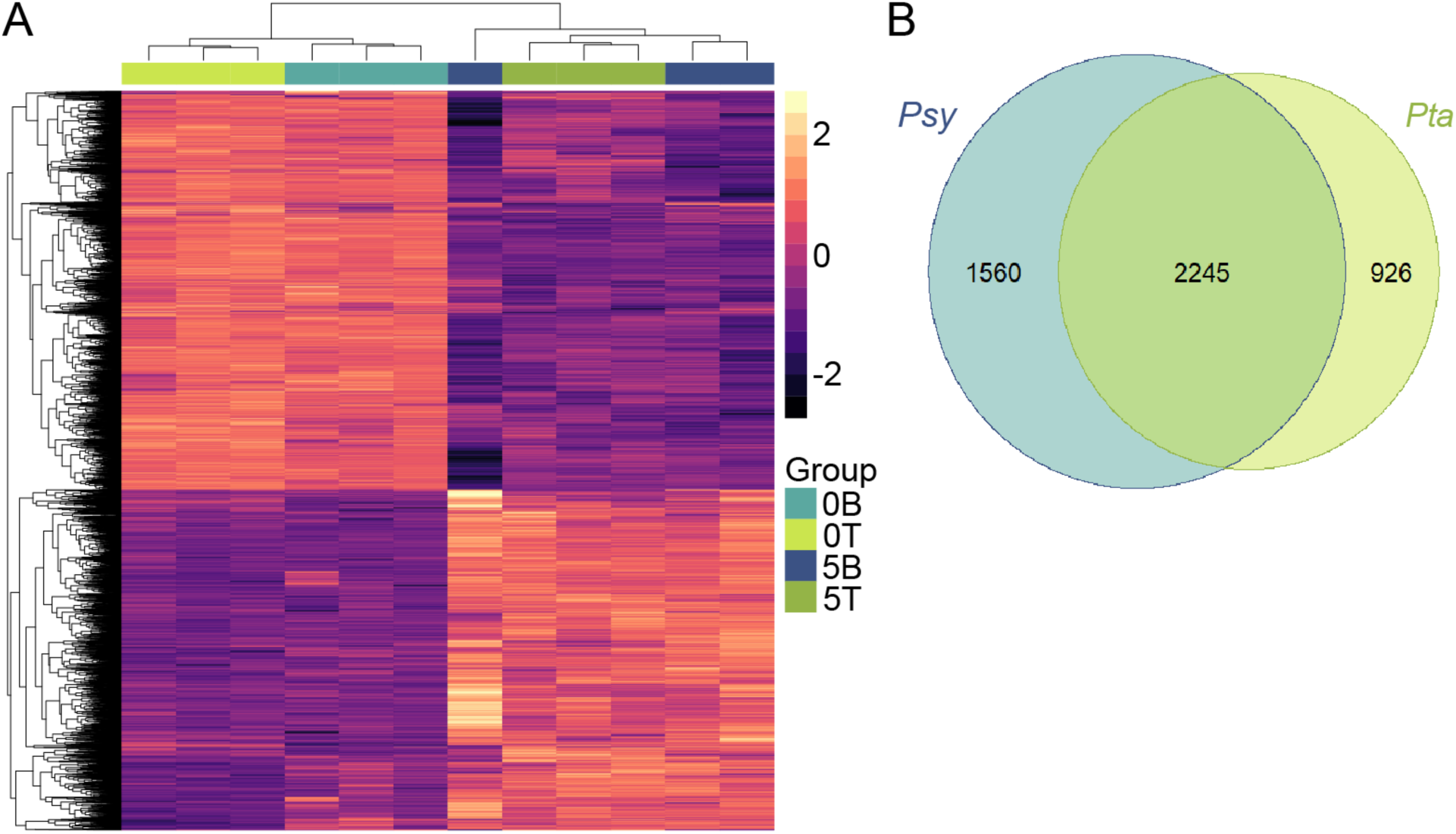
Differentially expressed genes in *Nicotiana benthamiana* in response to inoculation with *P. syringae* pv. *syringae* B728a (*Psy*) and *P. syringae* pv. *tabaci* ATCC11528 (*Pta*) five hours post inoculation compared to zero hours post inoculation. (A) Heat map showing regularized logarithm transformation of the count data for genes that are differentially expressed in response to *Pta* or *Psy* or both strains. 0B, uninoculated; 0T, uninoculated; 5B, 0B plants 5 hpi with *Psy*, 5T, 0T plants 5 hpi with *Pta*. (B) Venn Diagram showing total genes differentially expressed in response to *Pta* or *Psy* or both strains, when comparing 0 hpi to 5 hpi.

**Figure 2.**
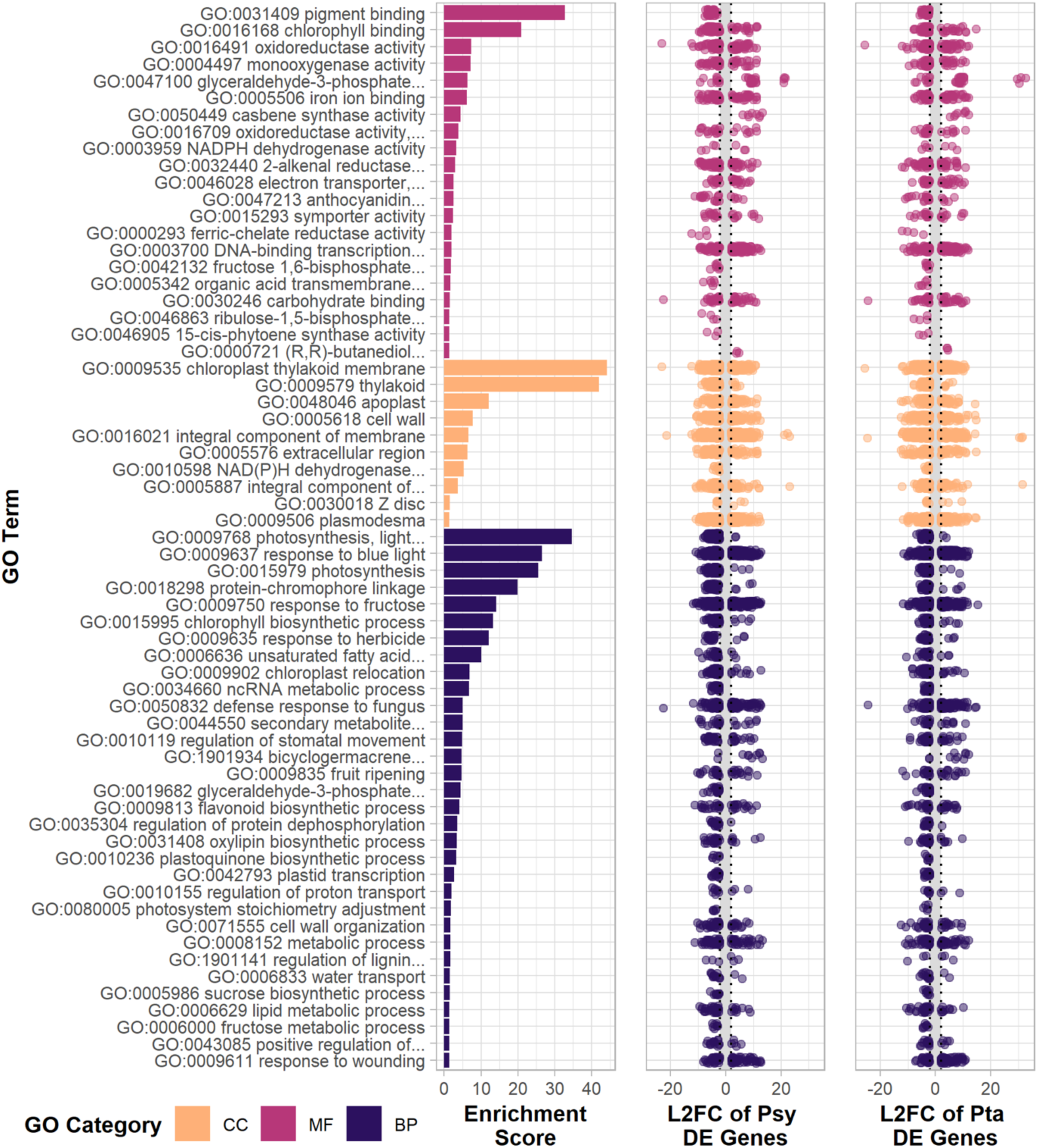
Gene ontology (GO) enrichment for differentially expressed (DE) genes shared in the responses of *Nicotiana benthamiana* to inoculation with *P. syringae* pv. *syringae* B728a (*Psy*) or *P. syringae* pv. *tabaci* ATCC11528 (*Pta*) at 5 hpi. GO terms have an adjusted p value<0.01 and were reduced in redundancy using Revigo. The full list of enriched GO terms is shown in Supplemental Figure 3. L2FC, log2 fold change; CC, cellular component; MF, molecular function; BP, biological processes.

We see defense response to fungus (GO:0050832) enriched in the shared DE genes (**Figure 2**), with associated genes mostly upregulated as expected by the introduction of a biotic stress. While the GO term appears specific to fungi, the overlap between immune recognition of pathogenic microbes among host organisms suggests genes within this term are not necessarily specific to plant-fungal defense. Other terms of interest related to known aspects of pathogenesis and defense include regulation of stomatal movement (GO:0010119), apoplast (GO:0048046), and oxylipin biosynthetic processes (GO:0031408); the plant hormone jasmonic acid and other oxylipins play a large role in plant development and stress response (Creelman and Mulpuri 2002). A PR1 ortholog (NbD052315) is highly downregulated in both (L2FC_Psy_=-11.65, L2FC_Pta_=-9.94). PR1 is widely used as an indicator of immune system activation and as a marker of salicylic acid immune signaling against biotrophic pathogens such as P. syringae (Han et al. 2023). The downregulation of PR1 is consistent with predicted immune system dampening by compatible phytopathogens.

### Plant response to Psy involves more chloroplast-related genes

We chose to focus the rest of our inquiry on the 1560 and 926 genes DE only in response to infection with *Psy* or *Pta*, respectively (**Figure 1B**, **Figure 3**), and we highlight that differential responses in these genes are not likely to be due to environmental causes other than infection. In response to *Psy*, 651 genes are upregulated and 909 are downregulated. Auxin signaling and manipulation plays an extensive role during plant-pathogen interactions (Kunkel and Harper 2018), and, of the ten DE genes with a L2FC>15 in response to *Psy* (**Table 1**), a relatively large number are involved in auxin related pathways. Three are related to auxin response and homeostasis (NbD025458, NbD037056, and NbD041542), while NbD025458, the highest upregulated, codes for a putative CHD3/CHD4-like chromatin remodeling protein. The second highest upregulated, NbD037056, is a BIG auxin transport protein. Eleven other upregulated genes are predicted IAA/Aux or auxin response factor (ARF) transcriptional activators or repressors. Overall, 60 upregulated genes have auxin-related GO terms; 46 auxin-related genes are downregulated, including some related to brassinosteroid and abscisic acid signaling such as orthologs of HAT1, PILS5, and TTL1. NbD041542 is a Protein DETOXIFICATION 48 homolog, a multidrug and toxin extrusion transporter that can mediate iron homeostasis under stress (Seo et al. 2012). An additional upregulated gene is a putative callose synthase (NbD005055), but only certain callose synthases are important for plant defense, with others involved in development and division and plasmodesmatal permeability control (Ellinger and Voigt 2014; Wu et al. 2018). NbD005055 was computationally predicted as callose synthase 1- or 2-like, which are both known to be induced are part of the defense response to pathogens in Arabidopsis in a salicylic acid-dependent manner (Dong et al. 2008), but are also described as serving roles in cell division (Ellinger and Voigt 2014). The four most downregulated genes are mostly uncharacterized, though one (NbD033797) has similarity to a soybean transcription factor and another (NbD000103) is related to a positive regulator of floral development in Arabidopsis (Murtas et al. 2003), which hints at a link between defense and development.

**Figure 3.**
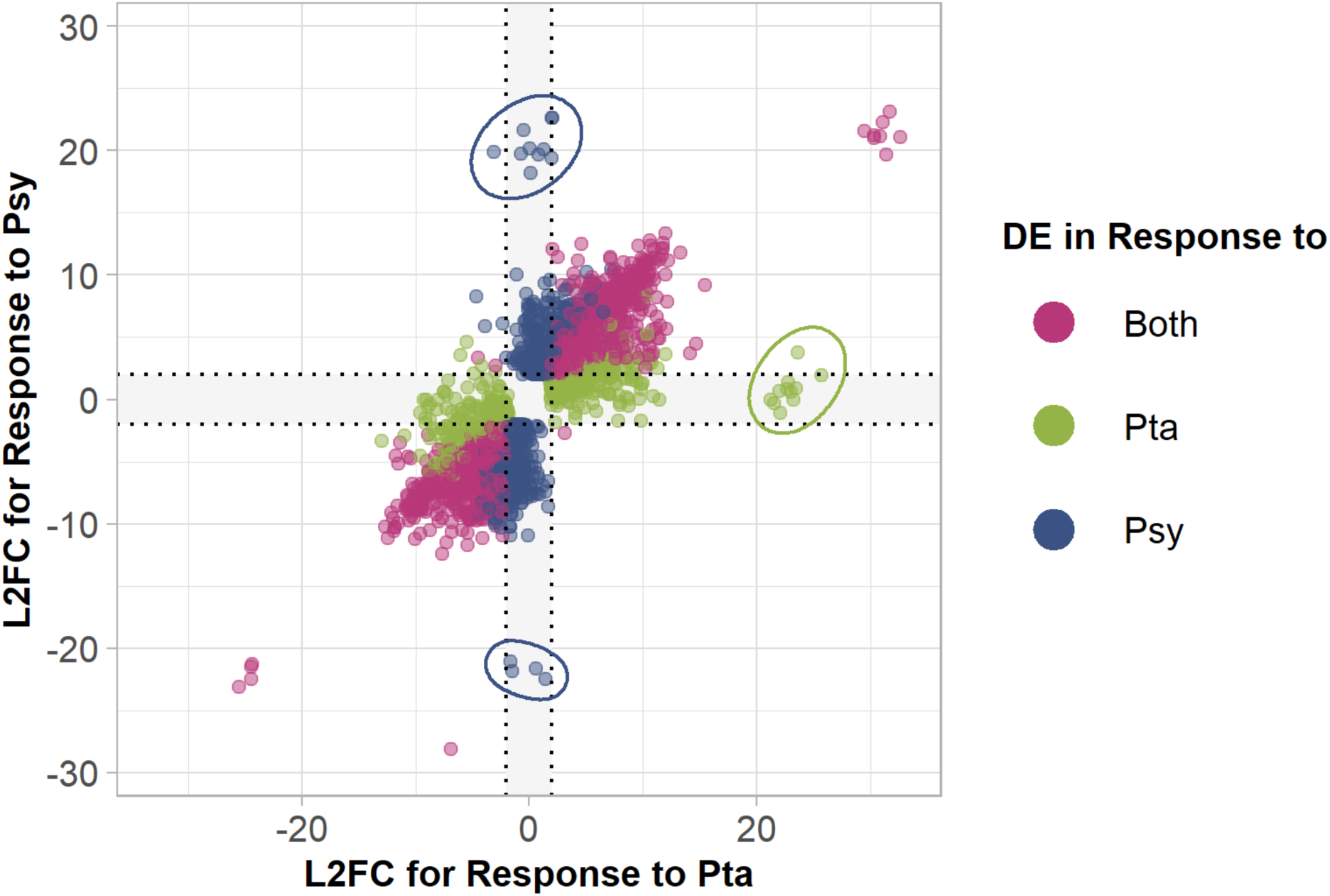
Scatterplot of the *Nicotiana benthamiana* genes that are differentially expressed (DE) in response to *P. syringae* pv. *syringae* B728a (*Psy*) or with *P. syringae* pv. *tabaci* ATCC11528 (*Pta),* or both. The axes represent that log_2_ fold change (L2FC) of the genes in response to one pathogen or the other. The circles indicate the most DE genes in response to one pathogen that are listed in **Tables 1 and 2**.

**Table 1.** Genes differentially expressed by *Nicotiana benthamiana* uniquely in response to *Pseudomonas syringae* pv*. syringae* B728a at 5 hours post inoculation compared to uninoculated with an absolute value of log2 fold change (L2FC) greater than 15.

| Gene | Product | Gene Name* | Description* | L2FC |
| --- | --- | --- | --- | --- |
| NbD025458 | uncharacterized protein<br>LOC104212182 | *chd4* | CHD3-type chromatin-remodeling factor PICKLE | 22.61 |
| NbD037056 | auxin transport protein<br>BIG isoform X1 | BIG | Auxin transport protein BIG | 22.60 |
| NbD010045 | ribosomal protein S13<br>(mitochondrion) | *rps13* | Ribosomal protein S13 | 21.65 |
| NbD005055 | callose synthase 2-like | CALS1 | Callose synthase 1 | 20.20 |
| NbD042013 | cleavage and<br>polyadenylation specificity<br>factor subunit 3-II | FEG | Integrator complex subunit 11 | 20.09 |
| NbD031078 | uncharacterized protein<br>LOC109215177 | At2g38970 | C3HC4-type RING finger-containing protein | 19.89 |
| NbD052852 | ubiquitin-conjugating<br>enzyme E2 variant 1C-like, partial | UEV1D | Ubiquitin-conjugating<br>enzyme E2 variant 1D | 19.76 |
| NbD026296 | MAP3K epsilon protein<br>kinase 1-like | M3KE1 | MAP3K epsilon protein<br>kinase 1 | 19.70 |
| NbD041542 | protein<br>DETOXIFICATION 48-like | DTX48 | Protein DETOXIFICATION 48 | 19.42 |
| NbD032348 | rho GTPase-activating<br>protein 7-like isoform X1 | ROPGAP7 | Rho GTPase-activating<br>protein 7 | 18.17 |
| NbD020455 | Partial, uncharacterized protein LOC109226839 | - | - | -21.06 |
| NbD000103 | Partial, ubiquitin-like-specific protease ESD4 | ESD4 | Ubiquitin-like-specific protease ESD4 | -21.68 |
| NbD019312 | uncharacterized protein LOC104238947 | At1g12380 | F5O11.10 | -21.88 |
| NbD033797 | uncharacterized protein LOC107824489 | glysoja_046371 | Transcription factor, putative | -22.46 |

\*UniRef90 BLAST top hit as annotated by Sma3s v2 (Casimiro-Soriguer, Muñoz-Mérida, and Pérez-Pulido 2017)

**Table 2.**
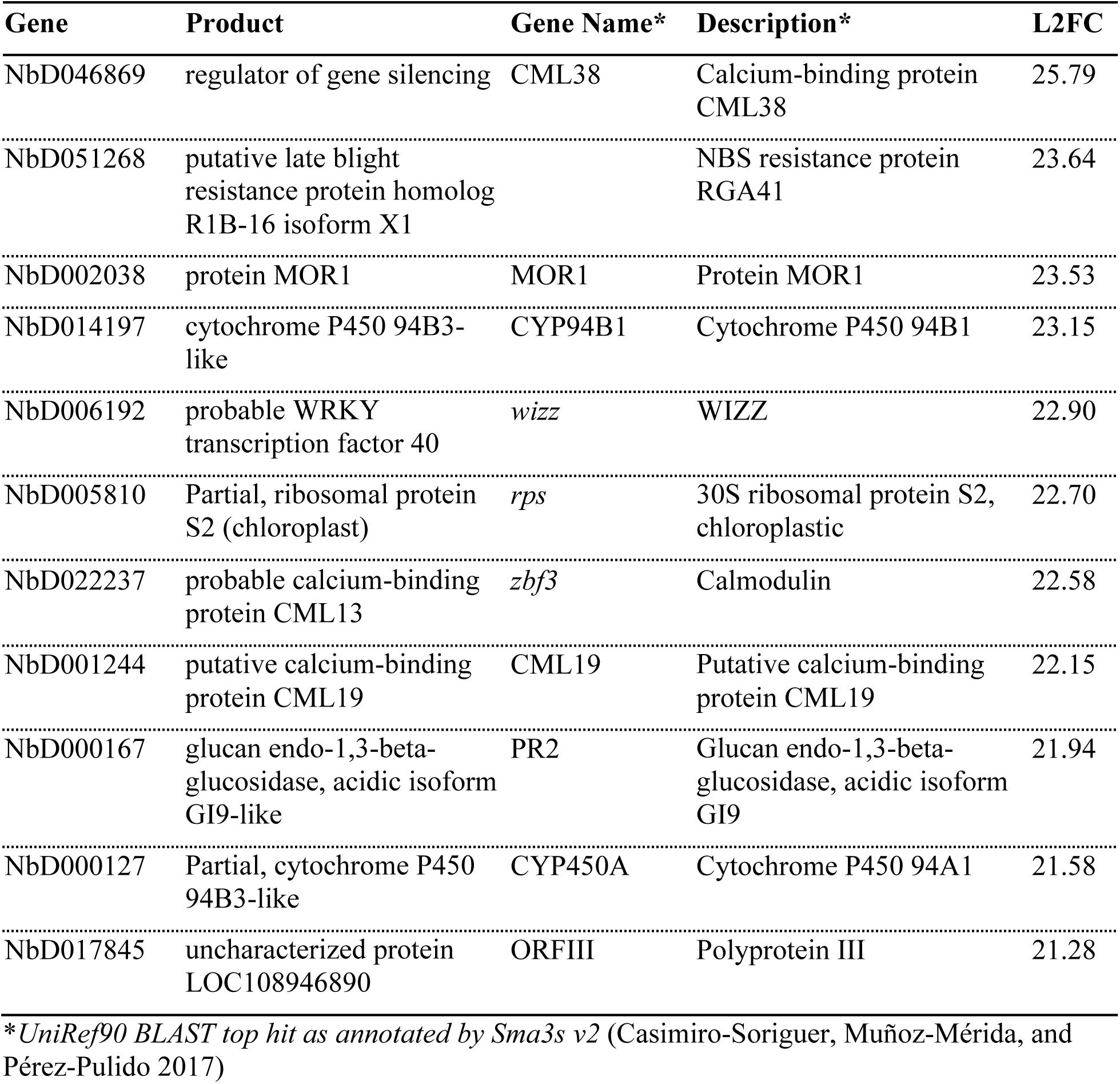
Genes differentially expressed by *Nicotiana benthamiana* uniquely in response to *Pseudomonas syringae* pv*. tabaci* ATCC11528 at 5 hours post inoculation compared to uninoculated with an absolute value of log2 fold change (L2FC) greater than 15.

| Gene | Product | Gene Name* | Description* | L2FC |
| --- | --- | --- | --- | --- |
| NbD046869 | regulator of gene silencing | CML38 | Calcium-binding protein CML38 | 25.79 |
| NbD051268 | putative late blight resistance protein homolog R1B-16 isoform X1 |  | NBS resistance protein RGA41 | 23.64 |
| NbD002038 | protein MOR1 | MOR1 | Protein MOR1 | 23.53 |
| NbD014197 | cytochrome P450 94B3-like | CYP94B1 | Cytochrome P450 94B1 | 23.15 |
| NbD006192 | probable WRKY transcription factor 40 | wizz | WIZZ | 22.90 |
| NbD005810 | Partial, ribosomal protein S2 (chloroplast) | rps | 30S ribosomal protein S2, chloroplastic | 22.70 |
| NbD022237 | probable calcium-binding protein CML13 | zbf3 | Calmodulin | 22.58 |
| NbD001244 | putative calcium-binding protein CML19 | CML19 | Putative calcium-binding protein CML19 | 22.15 |
| NbD000167 | glucan endo-1,3-beta-glucosidase, acidic isoform GI9-like | PR2 | Glucan endo-1,3-beta-glucosidase, acidic isoform GI9 | 21.94 |
| NbD000127 | Partial, cytochrome P450 94B3-like | CYP450A | Cytochrome P450 94A1 | 21.58 |
| NbD017845 | uncharacterized protein LOC108946890 | ORFIII | Polyprotein III | 21.28 |

\*UniRef90 BLAST top hit as annotated by Sma3s v2 (Casimiro-Soriguer, Muñoz-Mérida, and Pérez-Pulido 2017)

In a clear trend, genes DE in response to *Psy* were enriched for GO terms relating to chloroplasts (**Figure 4**), including the top two cellular component (CC) terms of chloroplast envelope (GO:0009941) and thylakoid membrane (GO: 0009579). Of the 1560 DE genes, 380 of them have enriched chloroplast-related CC GO terms and of those, 326 are downregulated. Four of the 54 upregulated genes are related to glutathione redox signaling; two glutathione S-transferases (NbD007461, NbD045751), glutathione reductase (gr;NbD022932), and glutathione peroxidase (gpx; NbD036458). The most upregulated gene with the chloroplast envelope GO term is a 9-divinyl ether synthase (DES; NbD030668); under biotic stress, related proteins can form antimicrobial oxylipins that have been found to be more useful against eukaryotic pathogens (Prost et al. 2005). For example, NtDES1 from *N. tabacum* is upregulated in the early response to a pathogenic oomycete, while other oxylipin synthesis genes have roles in defense against nematode and fungal infections (Fammartino et al. 2007; Wang et al. 2021; Sanadhya et al. 2021). In contrast, the most downregulated gene with the chloroplast envelope GO term is a receptor-like kinase (NbD047510) from the THESEUS 1/FERONIA family that can serve to detect cell wall defects and damage (Cheung and Wu 2011); this gene is annotated with additional GO terms associated with other cellular membranes, indicating unclear predicted localization.

**Figure 4.**
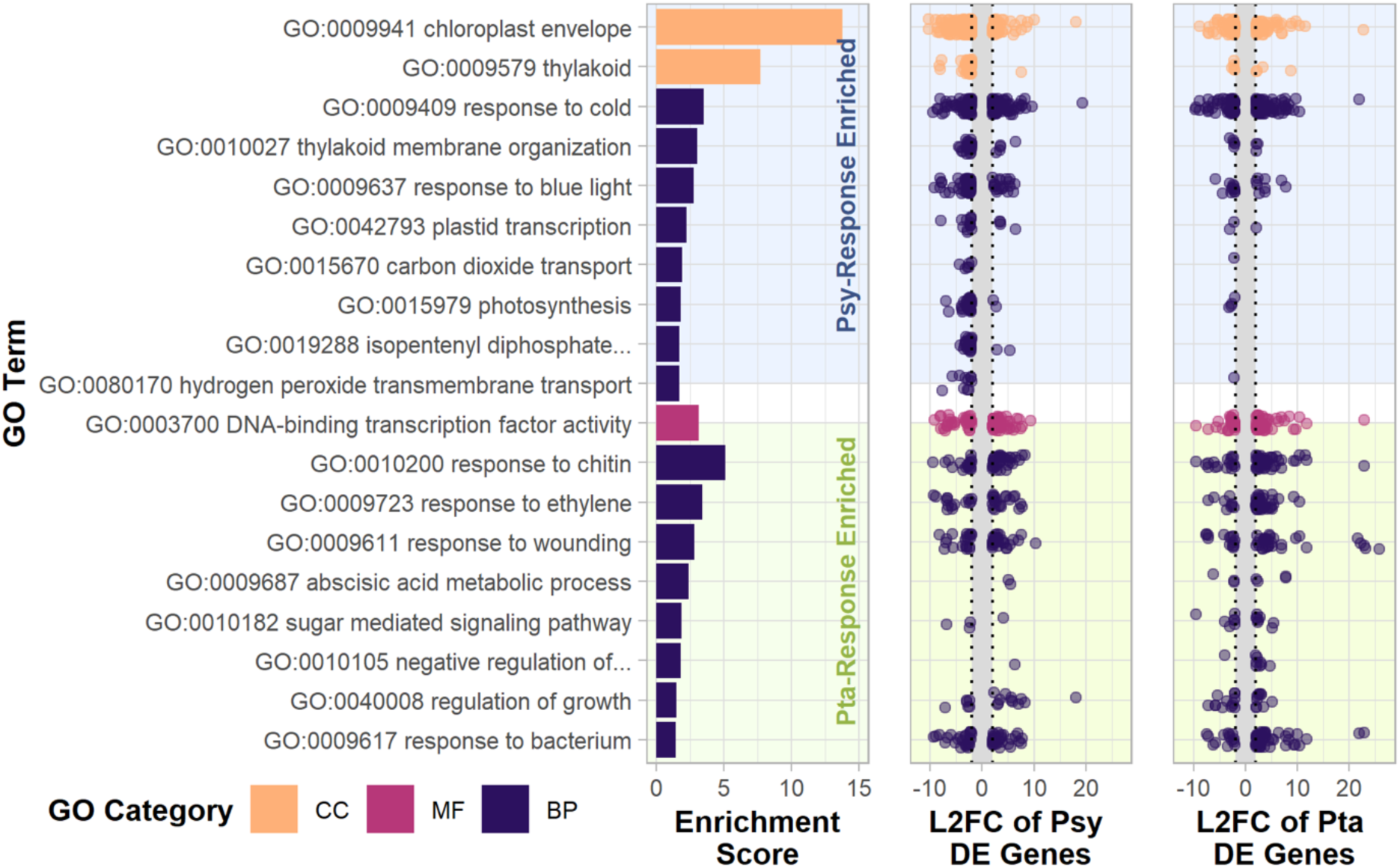
Gene ontology (GO) enrichment for differentially expressed (DE) genes specific to the response of *Nicotiana benthamiana* to inoculation with *P. syringae* pv. *syringae* B728a (*Psy*) or with *P. syringae* pv. *tabaci* ATCC11528 (*Pta)* at 5 hpi. GO terms were considered enriched for an adjusted *p* value<0.05. L2FC, log_2_ fold change; CC, cellular component; MF, molecular function; BP, biological processes.

### Plant response to Pta involves more hormone and calcium signaling

In total, 489 genes were upregulated and 437 downregulated specifically in response to *Pta* infection. Of the most DE genes (**Figure 3**), three are calcium binding proteins and two are cytochrome P450s (**Table 2**). Related cytochrome P450s CYP94B1, CYP95B3, and CYP94A1 regulate and interact with jasmonic acid signaling in Arabidopsis and *Vicia sativa* (Koo et al. 2014; Pinot et al. 1998; Heitz et al. 2012). Calcium signaling positively and negatively regulates the response to plant pathogens via calcium binding proteins like calmodulin (Zhang, Du, and Poovaiah 2014). For example, cellular calcium ion concentration changes can lead to negative regulation of the salicylic acid pathway by downregulating gene expression of EDS1 by AtSR1, dependent on calmodulin binding (Du et al. 2009). Virulent *P. syringae* pv. *tomato* can increase intracellular calcium, resulting in downregulation of salicylic acid receptor NPR1 by AtSR1, as well (Yuan, Tanaka, and Poovaiah 2021). Though, these calcium-binding proteins may be serving other roles, as an EDS1 homolog (NbD037498) and two NPR3 homologs (NbD049359, NbD051607) are upregulated by a L2FC of 2.7 to 4 in the shared response, or response to *Psy* for NbD051607. Another of the most DE genes is a nucleotide-binding leucine-rich repeat protein (NbD051268), a class of resistance proteins involved in pathogen perception and defense; this one must not confer resistance to these *Pseudomonas* strains given the compatibility for disease progression.

In contrast to the response to *Psy*, there are no enriched CC GO terms (**Figure 4**) in the DE genes specific to the *Pta* infection response. The majority of the enriched BP GO terms are for responses to biotic and abiotic stresses with the top three being response to chitin (GO:0010200), response to ethylene (GO:0009723), and response to wounding (GO:0009611). Of the 51 DE genes with ethylene-related enriched GO terms (GO:0009723, GO:0009873, GO:0010105), 41 are upregulated and 9 are downregulated. Among the upregulated genes are two putative RMA1H1-like E3 ubiquitin-protein ligases, which inhibit aquaporin trafficking during drought stress in hot pepper (Lee et al. 2009). Many upregulated ethylene response genes included transcription factors ilike orthologs of ERF1, ERF2, ERF3, ERF4, ERF5, ERF011, EIN3, EIN4, and RAV1. In fact, putative transcription factors (TF) are generally enriched in the upregulated genes in response to *Pta* as indicated by the molecular function (MF) GO term DNA-binding transcription factor (GO:0003700). Beyond the ethylene-response TFs, two orthologs of the wound-response TF WIZZ (Hara et al. 2000) are highly upregulated, followed by zinc-finger binding proteins and two orthologs of fungal-response TF WRKY33 (Zheng et al. 2006).

### Bacterial virulence genes typically followed expected expression patterns in planta

In parallel, we assessed the changes in gene expression within the bacteria, identifying orthologous pairs to compare and contrast their virulence strategies. *Psy* and *Pta* differentially express 462 orthologous genes (**Fig. 5**) five hours post infection *in planta* compared to growth on solid agar followed by resuspension and centrifugation. An additional 249 orthologous genes and 141 unique genes are DE in *Pta*, and 591 and 161 in *Psy*, respectively. The type III secretion system is one of the most important pathways involved for virulence *in planta* for *Pseudomonas syringae*, and many structural and regulatory components of this pathway are differentially expressed genes across both pathogens *in planta* (**Fig. 6**). Likewise, many effector proteins that are known to be translocated across the type III secretion system are also either differentially upregulated or trend in that direction for both *Psy* and *Pta*, including seven shared effector proteins (*avrE, hopI*, *hopAG1*, *hopAE1*, *hopAB1*, *hopX1*, and although truncated in *Pta*, *hopAA1*). Interestingly, we found that the core effector *hopM* was upregulated in *Pta* but not *Psy* and that there was variation in expression patterns for *hopAH* alleles across strains, with *hopAH2* in *Psy* trending towards upregulated *in planta.* This is of note because, although *hopAH2* populates many effector lists, the effector status of this locus has been long questioned given that it does not have a clearly identifiable *hrp* box in *Psy* although it is capable of being delivered via the T3SS. Most of the effector genes that are uniquely found in one pathogen but not the other are also either differentially expressed or trend that way with one interesting exception. *hop*W1 is found in four identical copies throughout the genome of *Pta*, including one copy present on a plasmid. Precise measurement of expression differences for these loci is difficult because of multimapping to identical sequences, but if one assumes equal expression across these *hopW1* loci, none of the loci individually is differentially expressed in a strong enough way to reach statistical significance *in planta* in *Pta*. However, that there is a significant upregulation of expression in *hopW1* if we map all reads at both time points to a single locus, suggests the potential for a higher level of transcription overall for *hopW1* during infection.

**Figure 5.**
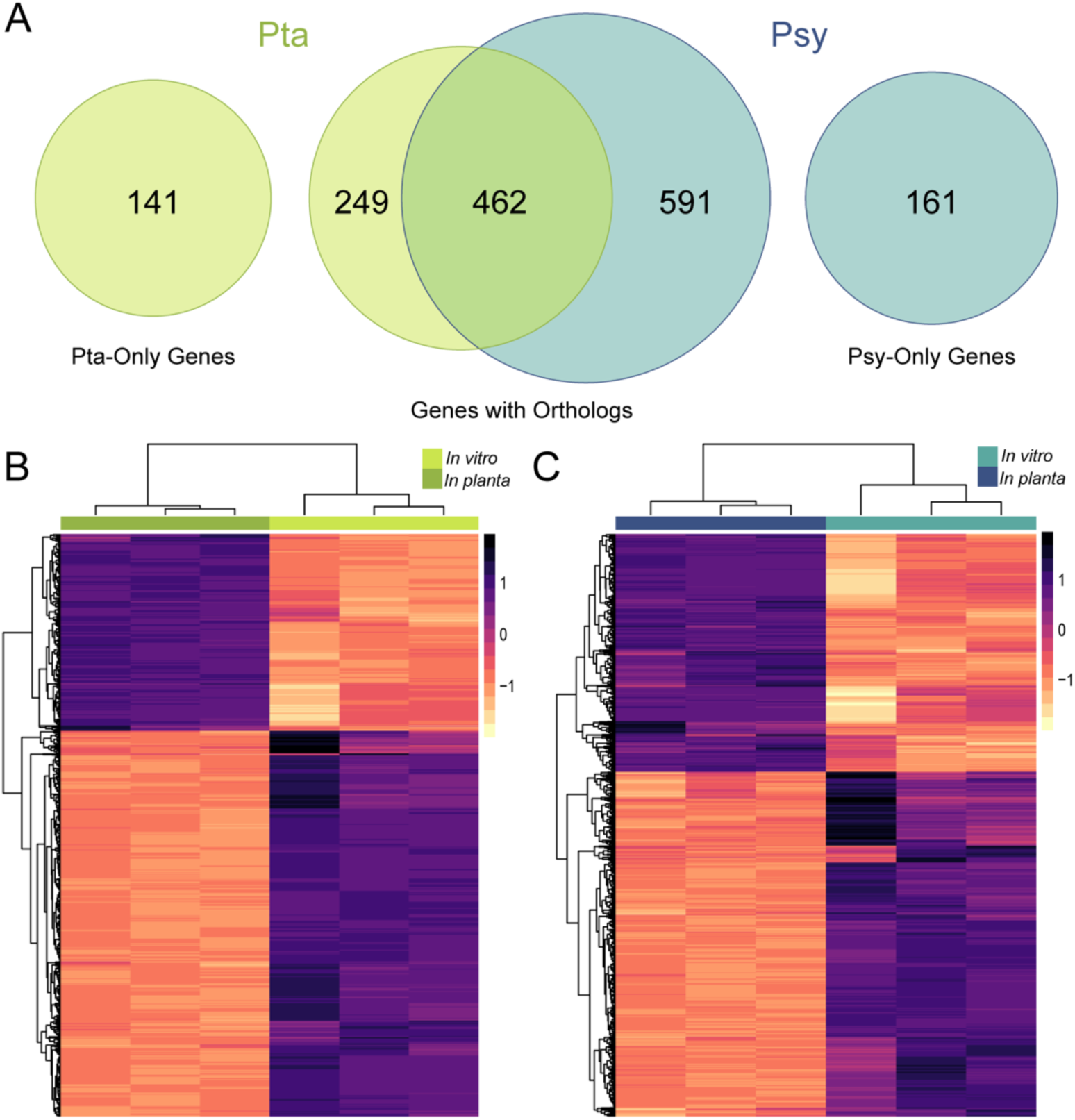
Differential gene expression in *Pseudomonas* strains (*P. syringae* pv. *syringae* B728a *Psy*; *P. syringae* pv. *tabaci* ATCC11528, *Pta) in vitro* compared to *in planta* five hours post inoculation. (A) Venn diagram depicting the number of differentially expressed genes with and without orthologs in *Pta* and *Psy*. (B-C) Heat map showing regularized logarithm transformation of the count data for genes that are differentially expressed by (B) *Pta* and (C) *Psy* during early plant infection.

**Figure 6.**
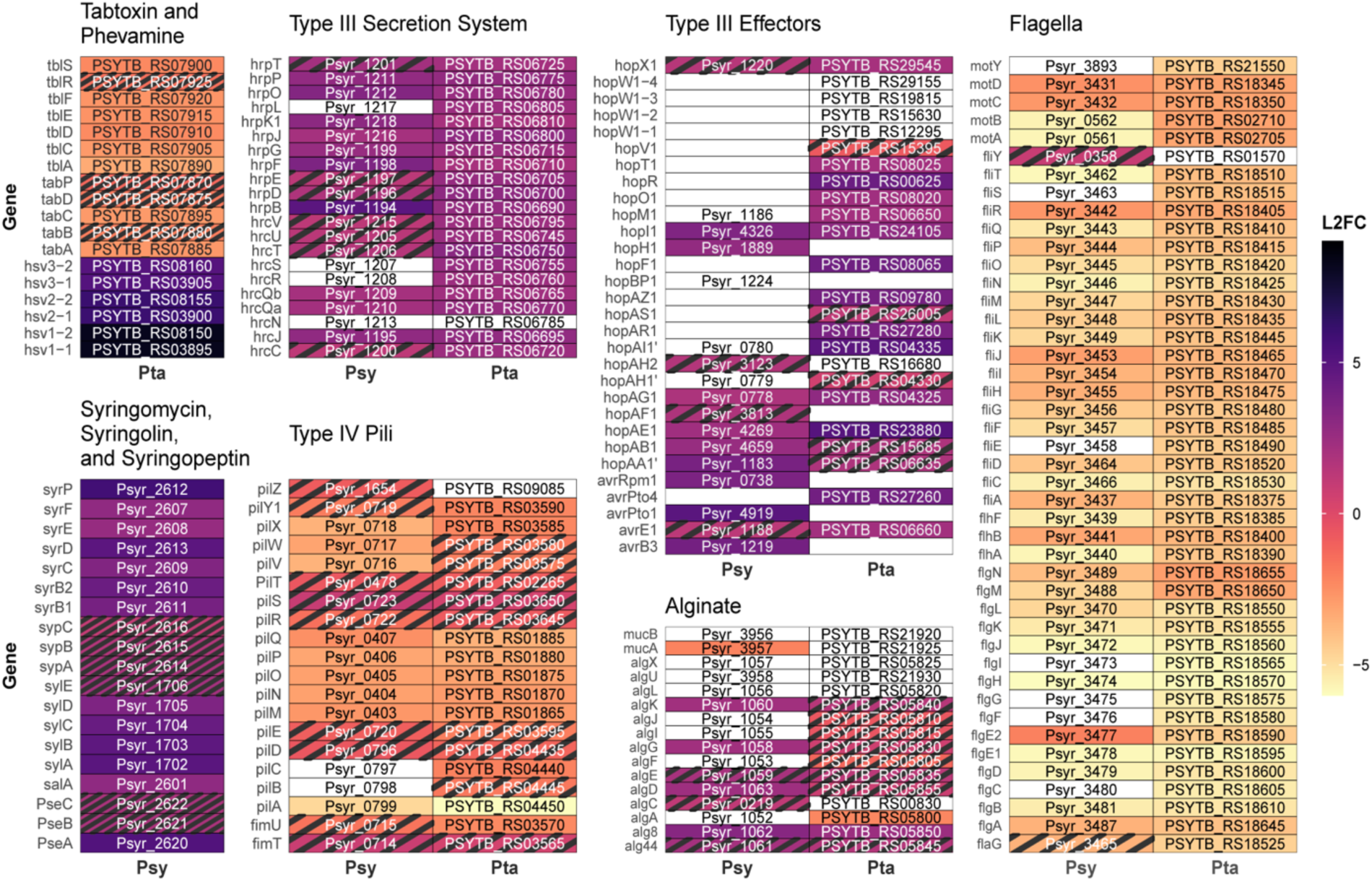
Differential expression of select virulence-related genes and pathways in *P. syringae* pv. *syringae* B728a (*Psy*) and *P. syringae* pv. *tabaci* ATCC11528 (*Pta) in planta* based on an adjusted p value<0.01 and |log2 fold change|>2 . Striped boxes indicate genes that are significant for different thresholds: adjusted p value<0.05 and no minimum log2 fold change (L2FC). White boxes indicate no ortholog present or, when labeled with a locus ID, orthologs that are not differentially expressed for either set of thresholds.

These two pathogenic strains do not share phytotoxin pathways, but we find that most of these pathways also appear to be largely upregulated *in planta* at 5hpi. For instance, all known phytotoxins in *Psy* are differentially upregulated *in planta* at 5 hpi. These include pathways for syringomycin production (*syr*B1, *syr*B2, *syr*CDEF), syringopeptin production (*syp*ABC) and syringolin production and export (*syl*ABCDE, *pse*ABC). Likewise, there are two copies of the operon predicted to encode phevamine A in *Pta*, and all of the genes in each (*hsv*ABC) are highly upregulated. Surprisingly, however, we find that loci involved in the regulation and production of tabtoxin (*tab*ABCDP, *tbl*ACDEFRS) are all significantly downregulated *in planta* compared to growth on solid agar. We note that loci involved in tabtoxin production have previously been found to be highly expressed during growth in culture under laboratory conditions (Barta et al. 1993), that genes for tabtoxin are still highly expressed *in planta* compared to other pathways (but just lower than under laboratory culture conditions), and that follow up experiments with qPCR from *in planta* samples confirmed downregulation of tabtoxin *in planta* compared to growth on agar plates (Supplemental Figure S7).

### Bacterial Expression Responses of Additional Genes *In Planta*

There is a dominant overall signal of growth differences for both strains *in planta* compared to on solid agar. This signal is represented as a variety of “housekeeping” genes involved in basic growth processes like cell division and translation being differentially expressed during infection, and somewhat obscures our ability to identify “newly” expressed genes *in planta* because these could likely just represent correlates of growth. However, we find that both strains respond in quite consistent and convergent ways *in planta* for a variety of pathways that could potentially impact virulence. Notably, genes involved in both flagellar and pili-based motility are downregulated in both *Pta* and *Psy* during infection while genes involved in alginate production trend towards upregulation in both (**Figure 6**). We note that follow up RT-qPCR results investigating *algD* expression *in planta* for *Pta* do suggest that RNAseq results for alginate production for this pathogen could be a lower estimate (Supplemental Fig. S7). Lastly, operons involved in the production of phage derived tailocin molecules are strongly upregulated in *Pta* (PSYTB_RS25445 to PSYTB_RS25575) as well as for *Psy* (Psyr_4582 to Psyr_4608) (**Supplemental Figure S5**). Since tailocin production is upregulated in response to DNA damaging agents under laboratory conditions, upregulation of tailocin pathways in both strains *in planta* could signal that this environment is challenging for bacteria even during “compatible” infections.

## Discussion

Extensive research over numerous decades has enabled fine scale characterization of the genetic basis of the variety of overlapping defense responses used by plants to defend against bacterial pathogens. Along the same lines, diverse arsenals used by various pathogens to overcome these defense responses have also been detailed using established model systems across plant hosts, with research highlighting the many ways that bacteria can disrupt plant host defenses. However, there has been relatively less emphasis on comparing and contrasting the genetic basis of compatible infections (where the outcome is disease) within a single host during infection by divergent phytopathogens, although responses are largely assumed to be correlated with overcoming common defense pathways.

Here, we report host responses during infection of *Nicotiana benthamiana* with two relatively closely related phytopathogens that are causative of different diseases in other hosts under natural conditions. Both pathogens are virulent in *N. benthamiana* when grown under laboratory conditions (Supplemental Figure S6), and we use this common host to test for underlying transcriptional convergence across pathogens in this host during the early stages of infection. While both pathogens (*Pseudomonas syringae* pv. syringae B728a, *Psy*; *Pseudomonas amygdali* pv. tabaci ATCC11528, *Pta*) fall under the banner of *Pseudomonas syringae* senso lato, they are classified as different phylogroups and species and potentially rely on different suites of virulence genes during compatible interactions (Baltrus et al. 2011). *Pta* is the causative agent of tobacco wildfire disease, and is thought to rely on a large number of type III effectors (18) as well as the tabtoxin and phevamine for growth and disease *in planta* (Baltrus et al. 2011; Studholme et al. 2009; O’Neill et al. 2018). *Psy* is the causative agent of brown spot of bean, and possesses fewer effectors than *Pta* (16) in addition to three different phytotoxins (syringomycin, syringolin, syringopeptin) (Feil et al. 2005; Helmann, Deutschbauer, and Lindow 2019; Vinatzer et al. 2006).

At a broad stroke, as reflected in Figs. 1, 2, and 3, plant responses to both pathogens are similar, with highly correlated levels of expression changes of the host plants to both pathogens. Notably, there are significantly different expression levels occurring in numerous pathways known to be involved in plant defense and both pathogens induce dramatic changes in chloroplast gene expression. We also note that, while it is possible that both pathogens are directly targeting and manipulating a subset or all of these pathways to cause disease, we cannot rule out that that these expression changes are also akin to a plant SOS message and are the product of general levels of stress as the internal plant environment degrades during infection. Despite perceived differences in virulence potential based on presence and absence of virulence factors across the *Psy* and *Pta* genomes, it does appear that infection by either pathogen tends to converge on similar changes within the host and that perceived differences in virulence strategies by the pathogen largely lead to similar outcomes, at least at this early interaction time point.

A subset of host responses significantly differs between the two pathogens, and these differences could signal more subtle divergence in virulence strategies between the pathogens themselves. Infection responses to *Psy* appear to lean more heavily on changes in chloroplast gene expression, while pathways that respond to *Pta* are biased towards those involved in response to regulation by plant hormones. As per Fig. 3, it is likely that all of these pathways generally respond in a similar way to both pathogens, but that effect sizes and variances across replicates of the responses are skewed in a way that yields statistical significance in only one of the two. It is also possible that these more subtle differences in response speak towards differences in the timing of infection. Since our sampling scheme had a standard time frame (5 hours post infection), it may be that both pathogens differ in when they affect common pathways and that these differences are being picked up in our larger analyses as strain specific responses. Indeed, we find that while both are ultimately virulent, that *Psy* displays a more rapid onset of symptoms and disease progression than *Pta* when grown under common laboratory conditions (Supplemental Figure S6).

Perhaps most interesting are genes in which expression changes significantly in response to one of the pathogens but which are virtually unchanged in the other pathogen (circled in Figure 3 and listed in Tables 1 and 2). Unlike many of the other genes that change significantly in response to infection in a correlated way between the two pathogens, and which therefore fall close to the diagonal in Figure 3, the circled genes are clearly biased in response to one of the pathogens. As above, these trends could reflect subtle timing differences between the pathogens during infection and thus might be significantly changed in both pathogens if measured at a different time scale. However, of any of the responses shown here, this class of genes is also the most likely to be responsive to only one of the pathogens and may reflect the outcomes of specific mechanisms of virulence employed by that strain. These genes also involve numerous loci that could be implicated in plant defense responses including pathways involved in callose production, ubiquitin ligases and proteases, MAP3K kinase and a WRKY transcription factor, calcium binding proteins, an NBS resistance protein, as well as a handful of uncharacterized proteins that will be interesting targets for future research efforts.

Prior to the experiments reported herein, and although we could make educated guesses based on previous research, it was unknown how similar (or different) a host plant’s responses to infection by two relatively closely related pathogens would be. We can now say that, at least for these two strains, that overall responses are broadly very similar across infection with these two pathogens at an early time point. Many of the significant differences are differences in magnitude but not in trend, and can potentially be explained by differences in the timing of disease development between these strains (Supplemental Figure S6). This convergence in host responses occurs despite divergence in the virulence arsenal between strains (both in terms of type III effectors and toxins). Conversely, there are a handful of genes that appear to be clearly and truly changed in regulation in response to one pathogen but not the other, and these genes are the best candidates for being the targets of strain specific virulence factors and potentially underlie or reflect the differences in disease phenotypes for these pathogens.

Comparison of gene expression signatures across these two bacterial strains offers a more complex story. The type III secretion system and effector proteins translocated by this system are thought critical for virulence of diverse *P. syringae* strains that infect diverse host plants. As expected, we find that many of the structural proteins involved in type III secretion and annotated type III effector proteins identified in each strain are upregulated at five hours post infection. These include effectors that are shared between both genomes as well as those that are exclusive to only one of the genomes. Expression of type III effectors is thought to be largely dependent on the action of the sigma factor HrpL, which binds to *hrp* boxes to recruit RNA polymerase to these regulons (Xiao and Hutcheson 1994; Ferreira et al. 2006; Lam et al. 2014). As part of this study, we have identified potential *hrp* boxes upstream of effector proteins and all effector loci identifed as upregulated *in planta* are located downstream of a potential *hrp* box promoter with the exception of Psy *hopBP1* (aka hopZ3). Both strains also contain phytotoxins that are demonstrated virulence factors. All three phytotoxins in *Psy* (syringomycin, syringopeptin, syringolin) are upregulated at 5 hpi compared to the controls. This suggests that, although toxins like syringolin have been shown to function to prevent stomatal closure to initiate infection (Schellenberg, Ramel, and Dudler 2010), they could also carry out numerous additional virulence functions. Likewise, the small molecule phevamine A has been shown to dampen plant immune responses to compatible pathogens (O’Neill et al. 2018). The phevamine A production operon is duplicated in *Pta*, but both copies appear to be relatively highly expressed in this pathogen *in planta*. However, in stark contrast to these other results, all genes implicated in tabtoxin production in *Pta* appear to be downregulated at 5 hpi *in planta* and this result has been confirmed using RT-qPCR (Supplemental Figure S7). Although, to our knowledge, there has been little investigation into tabtoxin pathway expression in planta, the tabtoxin result is surprising as production of this toxin does contribute to disease symptom progression. While there has only been limited investigation into the regulation of the tabtoxin production pathway, but one report does suggest that expression of these tabtoxin biosynthetic genes is relatively high under laboratory culture conditions (Barta et al. 1993). Further interrogation of data reported here demonstrates that pathways for tabtoxin production remain relatively highly expressed in planta compared to other genes (Table S3) but are lower than the baseline comparison of growth under laboratory conditions. Moreover, while it is possible that tabtoxin is a virulence factor that contributes during early stages of infection, it remains a possibility that this toxin contributes to late virulence in planta or only under certain environmental conditions or in certain hosts. Indeed, this line of thinking would be in line with a molecular version of the well-known “disease triangle” framework.

We also highlight that, since many of the predicted virulence genes are upregulated and correlated in expression between in both pathogens at the time point measured, that any differences in expression of other pathways represents a true divergence in expression profiles and is not simply a correlate of the environment that strains may be experiencing differently. To this point, although strain *Psy* has been demonstrated to systemically travel throughout *Nicotiana* plants during infection (Misas-Villamil, Kolodziejek, and van der Hoorn 2011), we find that all genes implicated in flagellar and pili based motility within this pathogen are largely downregulated at 5 hpi and that this response is shared with *Pta*. Likewise, these two strains display similar expression patterns with regards to alginate production, as this pathway is upregulated in *Psy* but unchanged to slightly upregulated in *Pta* at the same time point. Of note, follow up experiments with RT-qPCR *in planta* suggest that our RNAseq data for *algD* in *Pta* provides a low end estimate of expression of this pathway during infection (Supplemental Figure S7). While alginate production has been proposed to play a role in *P. syringae* tolerance to osmotic and oxidative stress in the apoplast (Chang Woo-Suk et al. 2007; Keith et al. 2003) as well as interfering with plant immune calcium signaling (Aslam et al. 2008; Scrase-Field and Knight 2003), alginate has been observed to make strain-variable contributions to virulence. In *Psy* alginate synthesis gene mutants had decreased apoplastic fitness in bean (Helmann, Deutschbauer, and Lindow 2019). We are not aware of any study directly examining the contributions of alginate to *Pta* virulence.

Taken together, our data paint a picture in which a host plant responds in largely overlapping ways to infection by two independently evolved but closely related compatible pathogens. At least for at 5 hpi and for these two bacterial strains, *N. benthamiana* does not appear to discriminate as judged by gene expression outputs, even though infection dynamics and longer-term symptoms differ between *Pta* and *Psy*. Likewise, even though both pathogens maintain somewhat independent suites of virulence genes, many of these predicted virulence loci are upregulated *in planta*. That host plant gene expression responses largely converge despite underlying differences in virulence factor repertoires likely speaks to both the redundancy of virulence factors as well as the importance of manipulation of system level nodes that pathogens use to overcome plant defense. Ultimately, while there may be many different ways that one can break an egg, eventually the output still converges on an egg getting broken.

## Data Availability

A complete genome assembly for *P. syringae* pv. *tabaci* 11528 genome can be found at Genbank accessions CP042804.1 and CP042805.1. Sequencing reads for Illumina (trimmed) and Nanopore (raw) for this assembly can be found at SRA accessions SRX6691661 and SRX6691662, respectively.

For transcriptome data, raw reads and processed count data can be accessed through GEO (accession GSE201377: For reviewer access, go to https://www.ncbi.nlm.nih.gov/geo/query/acc.cgi?acc=GSE201377 and enter token wjalwaskjrwxxup into the box).

Scripts for recreating figures and analyses can be found on Github at doi.org/10.5281/zenodo.6743815

Supplemental files 1 and 2, containing reannotation of type 3 effectors and phytotoxin genes for both strains referenced in this manuscript and the RT-qPCR data, respectively, can be found as supplemental files but are also available on FigShare at doi: 10.6084/m9.figshare.20151770.

## Limitations of this Study

We acknowledge that our experimental setup (inoculation of plant leaves by syringe infiltration) differs from the natural routes of infection by both pathogens, which invade through natural openings like the stomata. As such, it is certainly possible that incorporating a more natural route of infection could lead to increased divergence in host responses or in the particular genes that are directly manipulated by pathogen presence. Likewise, natural infection could change the timing of the infection responses to further exacerbate any host expression changes associated with infection after specific time points. We also acknowledge that our comparison group to *in planta* expression data consists of strains under laboratory growth on solid media, and that many housekeeping genes are also differentially upregulated compared to this condition *in planta*. We further note that, although three independent replicate infections were sampled on different days, that our data represent a single RNAseq experiment involving a single time point comparison involving a very highly concentrated bacterial inoculum. Therefore, while our results capture the regulatory status of bacteria and plants under very specific conditions, these likely reflect much more well controlled conditions than are found during infection in agricultural fields. Our RNAseq sample for bacteria also involves centrifugation of apoplastic fluid from infected leaves, and thus could be a biased against cells that remained in leaves despite centrifugation. We acknowledge the challenge of separating plant transcriptional responses to the pathogen from both wounding responses (given syringe infiltration) as well as circadian responses . It is possible that our interpretation results could be affected by this comparison, although we have been careful to limit our interpretations given these limitations throughout the manuscript. Lastly, we note that the read types (single vs. paired) and length differ between the *Psy* and *Pta* bacterial experiments. While our results and interpretations could be biased at some level by this difference, we believe that convergence in overall bacterial expression patterns argues against such biases significantly affecting our interpretations.

## Acknowledgments

## Acknowledgements

This material is based upon High Performance Computing (HPC) resources supported by the University of Arizona TRIF, UITS, and Research, Innovation, and Impact (RII) and maintained by the UArizona Research Technologies department and the Georgia Genomics and Bioinformatics Core at the University of Georgia. This work was supported in part by the University of Georgia Office of Research as well as the University of Georgia College of Agricultural and Environmental Sciences’ Research Office.

## Author Contributions

Conceptualization - DAB, BHK

Methodology - DAB, BHK, AS

Software - MEC

Formal Analysis - MEC

Investigation - DAB, BHK, AS

Resources - DAB, BHK

Data Curation – MEC, DAB

Writing – Original Draft Preparation - MEC, DAB, BHK

Writing – Review & Editing Preparation - MEC, DAB, BHK

Visualization - MEC

Supervision - DAB, BHK

Project Administration - DAB, BHK

Funding Acquisition- BHK

## Materials and Methods

### Genome Sequencing and Assembly of Pta

For each genomic DNA extraction used in the assemblies reported here, a sample of this frozen stock was streaked to King’s Media B (KB) agar plates and single colonies were transferred to 2-mL of KB broth and grown overnight at 27°C in a shaking incubator at 220 rpm after which genomic DNA was isolated. Genomic DNA used for Illumina sequencing was isolated from a 2-mL overnight culture via the Promega (Madison, WI) Wizard kit with the manufacturer’s protocols. Genomic DNA for Nanopore sequencing was prepared independently using a Circulomics (Baltimore, MD) Nanobind High Molecular Weight DNA extraction kit. RNAse was added as per manufacturer’s protocols for all of the genomic isolations where specified.

Genomic DNA was sequenced by SNPsaurus (Eugene, OR) using an Illumina HiSeq 4000 instrument and following their standard workflow for library preparation and read trimming. This workflow uses a Nextera tagmentation kit for library generation, followed by sequencing that generated 150-bp paired-end reads, followed by trimming of adaptors from reads with the computational suite BBDuk (BBMap version 38.41) (9). This workflow generated a total of 2,087,458 trimmed paired reads and 630 Mbp of sequence (∼95x coverage). Genomic DNA was also sequenced by the Baltrus lab via an Oxford Nanopore MinION using a R9.4 flowcell, with 1 µg of DNA prepared using the LSK-109 kit without shearing or size selecting (other than using the Long Fragment buffer supplied with kit). Reads were called during sequencing using Guppy version 3.2.6 using a MinIT (ont-minit-release 19.10.3) for processing. Nanopore sequencing generated 101,281 reads with an N50 of 6,116 bp. Sequencing reads for Illumina (trimmed) and Nanopore (raw) can be found at SRA accessions SRX6691661 and SRX6691662, respectively.

Hybrid assembly of all read types was performed using Unicycler version 0.4.8 (Wick et al. 2017). Assembly resulted in a single chromosome: 6,133,558 bp and a single 68,162 bp plasmid. Both replicons were determined to be circular, and both were rotated per the Unicycler pipeline (8). The genome was annotated using the NCBI PGAP pipeline (Tatusova et al. 2016). Default parameters were used for all software. The assembled chromosome sequence can be found at accession CP042804.1 and the sequence of plasmid pTab1 can be found at CP042805.1.

### Inoculation and sample collection for RNAseq

Two-week soil germinated *Nicotiana benthamiana* seedlings were transplanted into 4- inch pots with SunGro 3B professional growing mix and fertilized with Peters Professional 20-20-20 water-soluble fertilizer at 1g/L of water. Seedlings we grown in a Conviron Adaptis growth chamber with 12 hours light (125 μmol/m^2^/s) at 26°C and 12 hours dark at 23°C for 4 weeks and then transferred to the laboratory plant growth room with 12 hours light (35 μmol/m^2^/s) and ambient room temperature. Plants were 6-13 weeks post-seeding at the time of use.

We carried out three independent replicate *in planta* infection experiments per bacterial strain in order to sample RNA, and have included a schematic of our workflow for a typical infection experiment as Supplemental Figure S1. For each infection experiment, samples were collected from two consecutive expanded leaves on each of two plants of the same age. Immediately prior to bacterial inoculation (T0 plant), four 4 mm diameter leaf discs were collected using disposable biopsy punches from each leaf for a total of 16 leaf punches correlating to approximately 2 cm^2^ leaf tissue and immediately frozen in Liquid Nitrogen (LN_2_) until processing. *Pta* and *Psy* inoculum was prepared directly from fresh KB plate cultures incubated at 28°C (King’s B; per 1 L = 20.0 g proteose peptone 3, 0.4 g MgSO_4_⋅7H_2_O, glycerol 10-mL, 2.0 g K_2_HPO_4_⋅3H_2_O, 18 g agar), resuspended in 0.25 mM sterile MgCl_2_ and standardized to OD_600_ 0.5. A 1-mL sample of bacterial inoculum (T0 bacteria) was retained and the bacteria were collected by centrifugation and flash frozen in LN_2_ until processing. Four leaves (two each on two plants, which had been previously sampled for T0 plant data points) were fully infiltrated with the cell suspensions using a 1-cc blunt syringe. At 5 hours post-inoculation (T5 plant), a corresponding treatment sample of leaf tissue was collected with biopsy punches as described for T0.

To physically separate the *P. syringae* bacteria from the leaf tissue at 5 hours post inoculation (T5 bacteria), we adapted the procedure used by Lovelace et al 2018 for use with *N. benthamiana* leaves. Briefly two *N. benthamian*a leaves each were detached and gently rolled up and inserted into the barrels of two 20-mL syringes. An RNA stabilizing buffer (De Wit et al. 2012), pH 5.2, was poured into each syringe, which was sealed and vacuum infiltrated at 95 kPa for 2 min, followed by a slow release of the vacuum. Vacuum-infiltration with RNA-stabilizing buffer was conducted twice on inoculated leaves. Excess RNA-stabilizing buffer was decanted and the syringes were placed into 50-mL conical tubes and were centrifuged at 1,000 × *g* for 10 min at 4°C to recover the intercellular wash fluid (O’Leary et al. 2014). The flow-through was concentrated by syringe filtration using a 0.20-μm Micropore Express Plus membrane placed within a removable filtering syringe tip adapter (Millipore, Billerica, MA, U.S.A.). Filters were removed from the adapter and immediately frozen in LN_2_ until processing.

Each of the original T0 bacterial samples for the three replicates from strain *Psy* were rendered unusable for downstream analyses due to an abnormally high proportion of rRNA reads within these libraries. Therefore, we carried out a separate set of three replicate RNAseq experiments (using bacterial populations resuspended from independently growth on agar plates) to approximate T0 bacteria points for comparison to the *Psy* T5 *in planta* samples. Similar steps were followed as described above, except that paired RNA samples were also collected for this strain prior to pelleting by centrifugation for comparison of gene expression for *Psy*B728a pre and post centrifugation. A comparison of gene expression patterns for non-pelleted T0, pelleted T0, and T5 samples is included as Supplemental Figure S8.

### RNA Extraction and Sequencing

Frozen samples were homogenized in 2-mL homogenization tubes with high-density zirconium beads (Glen Mills) using a Geno/Grinder (SPEX SamplePrep) for 1 min at 1,750 Hz with a LN_2_ chilled sample holder. RNA was extracted using the DirectZol RNA miniprep kit (Zymo Research). RNA samples were DNAse treated with the Turbo DNAse (Thermo-Fisher) followed by cleanup with the Monarch RNA Cleanup Kit (NEB). Reagents were used according to the manufacturer’s recommendations.

Plant mRNA sequencing libraries were prepared with the Illumina-compatible KAPA Stranded mRNA-seq Kit (Roche) by the Georgia Genomics and Bioinformatics Core (GGBC). The *Pta* RNA samples were rRNA depleted using both RiboZero Plant and RiboZero Bacteria (Illumina) in combination and RNA sequencing libraries were prepared in-house using the TruSeq Stranded Total RNA kit (Illumina). The *Psy* T5 RNA samples were rRNA depleted using the RiboMinus Bacterial kit (Thermo Fisher) and libraries were prepared with the Illumina-compatible KAPA Stranded RNA-seq Kit (Roche) by the GGBC. Library prep kits were used according to the manufacturer’s recommendations. Indexed libraries were pooled at ratios to target >5 million reads for bacterial RNA samples and 10-20 million reads for plant RNA samples and sequenced at the GGBC. The *Psy* T5 bacterial RNAseq libraries and *Psy-*inoculated Plant mRNAseq samples libraries were pooled and paired-end 75-bp reads were sequenced using the Illumina Nextseq 500 in High-output model. For the pooled *Pta*-inoculated plant RNA samples, paired-end 75-bp reads were sequenced using the Illumina Nextseq 500 in Mid-output model. For the pooled *Pta* bacteria RNA samples, single-end 75-bp reads were sequenced using the Illumina Nextseq 500 in High-output mode. *Psy* T0 bacteria samples (non-pelleted and pelleted) were prepared and sequenced by SeqCenter (Pittsburgh, PA) according to their typical workflow. These *Psy* T0 samples were DNAse treated with Invitrogen DNAse (RNAse free). Library preparation was performed using Illumina’s Stranded Total RNA Prep Ligation with Ribo-Zero Plus kit and 10 bp IDT for Illumina indices. Sequencing was done on a NextSeq2000 giving 2x51 bp reads. Demultiplexing, quality control, and adapter trimming was performed with bcl-convert (Illumina, v3.9.3) .

### RNA-Seq Analysis

For plant reads, we aligned to the *Nicotiana benthamiana* draft genome sequence v1.0.1 with improved NbD transcriptome (Bombarely et al. 2012; Kourelis et al. 2019) using HISAT2 v2.2.1 and then gene counts were assessed with StringTie v2.2.0 restricting the counts to known transcripts (Pertea et al. 2016). For bacterial reads, bowtie2 v2.4.1 was used for alignment to the *P. syringae* B728a (Genbank: CP000075.1) or *P. amygdali* pv. *tabaci* (Genbank: NZ_CP042804.1 and NZ_CP042805.1) genome and featureCounts from the subread package for gene counts, allowing for overlapping and multimapping partial counts. The sequencing reads from the *in vitro* sample of *Psy* taken at the time of the infection experiments contained too much ribosomal RNA and were not informative enough for differential expression analysis. Thus, we carried out independent RNAseq sampling of three additional replicates of *Psy* resuspended under the same conditions as *Pta* T0 samples. We used OMA standalone to determine orthologs between the *Psy* and *Pta* genomes (Altenhoff et al. 2019).

Raw reads and processed count data can be accessed through GEO (accession GSE201377: For reviewer access, go to https://www.ncbi.nlm.nih.gov/geo/query/acc.cgi?acc=GSE201377 and enter token wjalwaskjrwxxup into the box)

We used the R package DESeq2 v1.3.1 (Love, Huber, and Anders 2014) to identify differentially expressed genes based on an adjusted p value of less than 0.01 and an absolute value of log_2_ fold change greater than two. As part of the calculation, the threshold on Cook’s distance is typically set at 99% quantile of the F(p, m-p) distribution, where p is the number of coefficients being fitted and m is the number of samples according to the DESeq2 manual, but was changed to a 97% quantile for the plant counts to remove additional genes with clear outlier counts among condition replicates. Goseq v3.14 was used for gene ontology enrichment analysis accounting for transcript length bias (Young et al. 2010). Output from GOSeq was reduced for term redundancy with Revigo set to small (0.5) and the closest available relative species *Solanum lycopersicum* (Supek et al. 2011). Visualization was done with ggplot2 v3.3.5, VennDiagram v1.7.1, viridis v0.6.2, and pheatmap 1.0.12 in R v4.1.2.

### RT-qPCR Analysis

To confirm there was no overt transcriptional bias on *Pta* physically separated from plant tissues, we conducted RT-qPCR on total RNA isolated from whole, infected tissue. Plants were infiltrated with inoculum as previously, and 1.5 mg tissue was obtained by biopsy punch at 5 hpi. These samples were RNA-extracted as previously, with the exception that DNAse treatment was done using the kit’s provided DNAse on-column. 1 μ g RNA was converted to cDNA using QScript cDNA SuperMix (QuantaBio, Beverly MA, USA) according to the manufacturer’s instructions, resulting in a mixed pool of both plant and bacterial RNAs. Resulting cDNA was diluted in sterile water prior to further work to 10 ng/μl and stored at - 20 C.

All qPCRs were carried out on a StepOnePlus Real Time PCR System (Applied Biosystems, Waltham MA, USA), using Luna Universal Sybr Green master mix (NEB, Ipswich MA, USA) and 0.25uM each primer (Eurofins, Louisville KY, USA). Melt curve steps were included to confirm amplification of expected targets. All primers were designed from Genbank accession genomic data for *Pta* ATCC11528 using Geneious 10.2.6 (BioMatters, Boston MA, USA).

To differentiate and quantify bacterial rRNA among host plant RNAs, cDNA samples were first quantified by qPCR, using a *P. syringae*-specific 16S rRNA gblock (IDT, Coralville IA, USA) five-fold serial-diluted in water to generate a standard curve from 1.02×10^-6^ – 0.08 ng per reaction. Samples were diluted to 1 ng/μ l, then five-fold serial-diluted three-to-five times to target appropriate Cqs. RNA that had not been converted to cDNA was simultaneously tested to confirm that the reverse-transcription was not based on carried-over genomic DNA. These no-RT samples were found to be sufficiently cleared of gDNA. The samples were then normalized according to these quantitation results to the lowest-concentrated sample, by diluting working cDNA in sterile water. Normalized samples were then tested against four genes of interest: *algD*, *fliC*, *hrpL*, and *tblA*; *rpoD* was used as a reference gene. Output qPCR data were run through LinRegPCR (Ruijter et al. 2009) software (HFRC, Amsterdam, NL) to calculate real efficiencies and Cqs of samples, which in turn were used to calculate fold-change of the four GOIs using a modified “ddCt” (Livak) method in Excel (Microsoft, Redmond WA, USA).

16S rRNA:

Locus: NZ_CP042804.1:701180-702718 PSYTB_RS03350

qrt-16S-Ptab-F ACGGGTACTTGTACCTGGTGG

qrt-16S-Ptab-R CGTTTCCGAGCGTTATCCC

*algD*:

Locus:NZ_CP042804.1:c1201694-1200378 PSYTB_RS05855

qrt-*algD*-Ptab-f GAAGAACGGCGATCTGGAACTC

qrt-*algD*-Ptab-R CAGTACTGCGAACAACGATGG

*fliC*:

Locus: NZ_CP042804.1:c4015455-4014607 PSYTB_RS18530

qrt-*fliC*-Ptab-F TACCAACCTGAACGGCAAGAA

qrt-*fliC*-Ptab-R GCGCTCAGAGTCAGAGTGAT

*hrpL*: (Moreno-Pérez, Ramos, and Rodríguez-Moreno 2021)

Locus: NZ_CP042804.1:c123740-123264 PSYTB_RS00635 hrpL

qrt-*hrpL*-Ptab-F CCTAGTGATCCTTGATGC

qrt-*hrpL*-Ptab-r CAAGCAATCAATGGCTGC

*rpoD*: (González-Lamothe et al. 2012)

Locus: NZ_CP042804.1:5499167-5501017 PSYTB_RS25745 rpoD

qrt-*rpoD*-Ptab-f TGGGTCGTGAGCAGAAG

qrt-*rpoD*-Ptab-R CGGATGATGTCTTCCACC

*tblA:*

Locus: NZ_CP042804.1:c1622777-1622082 PSYTB_RS07890 tblA

qrt-Ptab-*tblA*-f GGCGATCTGGTGCAAGCCAA

qrt-Ptab-*tblA*-R CGGCTGTGTACGCATCCAC

## Supplemental

**Supplemental File 1** (excel file). Annotation of major virulence gene loci from strains *P. syringae* pv. *syringae* B728a (*Psy*) or *P. syringae* pv. *tabaci* ATCC11528 (*Pta*). Virulence genes for each strain are listed in tabs in the excel file, with separate tabs for type 3 effectors and for phytotoxins. Each sheet is split into sections for either *Psy* or *Pta*. Gene names and loci numbers are provided for genome annotation versions listed in the manuscript as well as nucleotide and protein sequences. For type 3 effectors, we further list whether a *hrp* box has been identified and include a sequence for potential *hrp* boxes associated with each locus (consensus sequence GGAACC(N15/N16/CCAC).

**Supplemental File 2** (excel file). Data and calculations from RT-qPCR experiments.

**Supplemental Table 1** (excel file). Output from differential expression analysis for DESeq2 for *Nicotiana benthamiana* genes in response to inoculation with *P. syringae* pv. *syringae* B728a (*Psy*) or *P. syringae* pv. *tabaci* ATCC11528 (*Pta*).

**Supplemental Table 2** (excel file). Output from differential expression analysis for DESeq2 for *P. syringae* pv. *syringae* B728a (*Psy*) contrasting *in vitro* and *in planta* conditions.

**Supplemental Table 3** (excel file). Output from differential expression analysis for DESeq2 for *P. syringae* pv. *tabaci* ATCC11528 (*Pta*) contrasting *in vitro* and *in planta* conditions.

**Supplemental Table 4** (excel file). Output from OMA indicating orthologs between *P. syringae* pv. *syringae* B728a (*Psy*) and *P. syringae* pv. *tabaci* ATCC11528 (*Pta*) genomes.

**Supplemental Table 5** (excel file). We have collected annotation information for coding potential for type three effector genes as well as phytotoxin pathways within the genomes of *P. syringae* pv. *syringae* B728a and *P. syringae* pv. *tabaci* ATCC11528. We include this information as two tabs within the linked spreadsheet.

**Supplemental Figure 1.**
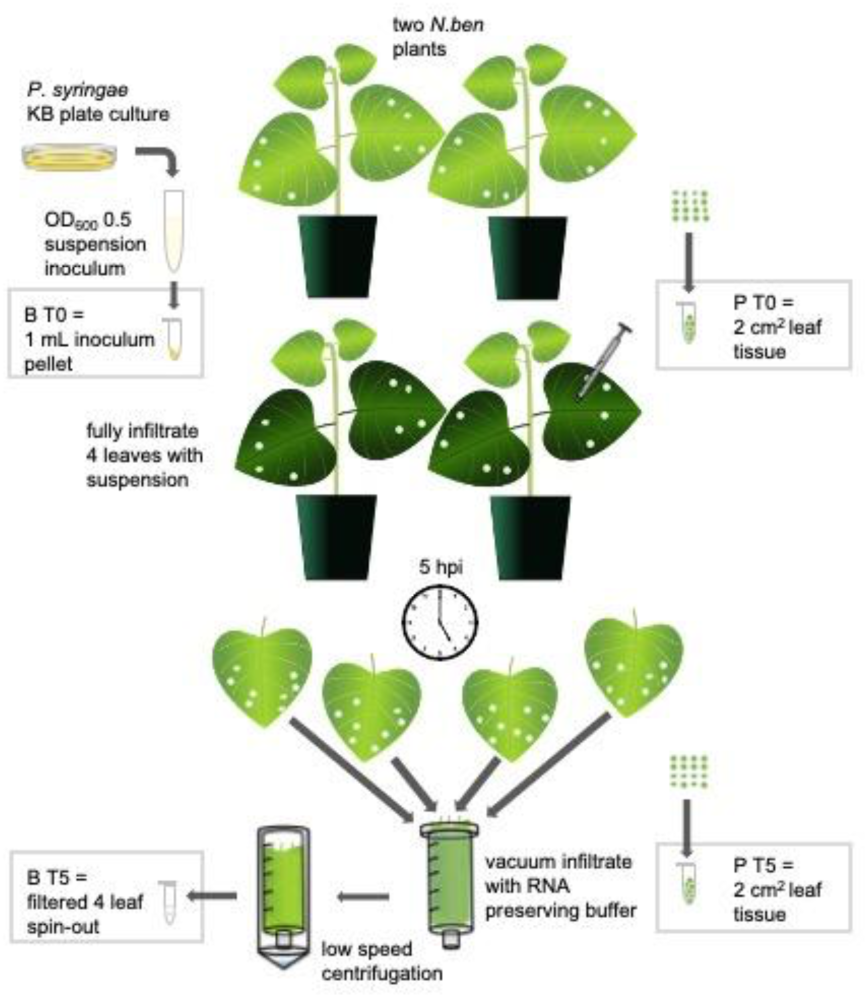
RNAseq Sampling Schematic. Procedures for a single biological replicate are shown. Experiments reported in this manuscript are the produce of three biological replicate infections per strain.

**Supplemental Figure 2.**
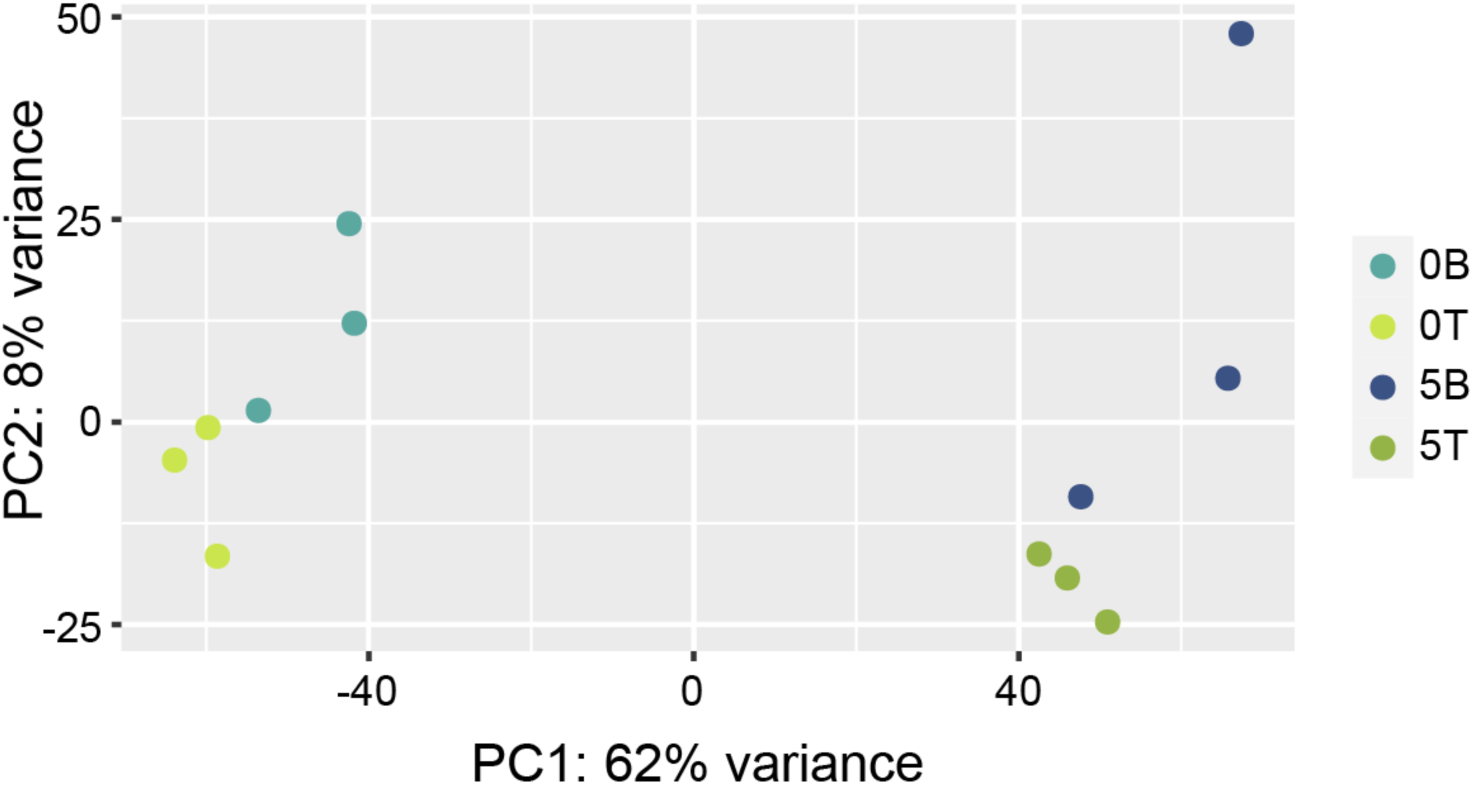
A principal component analysis plot depicting the three biological replicates for each of the four conditions sampled from *Nicotiana benthamiana* in the RNA-seq analysis. 0B, uninoculated; 0T, uninoculated; 5B, 0B plants 5 hpi with *Psy*, 5T, 0T plants 5 hpi with *Pta*.

**Supplemental Figure 3.**
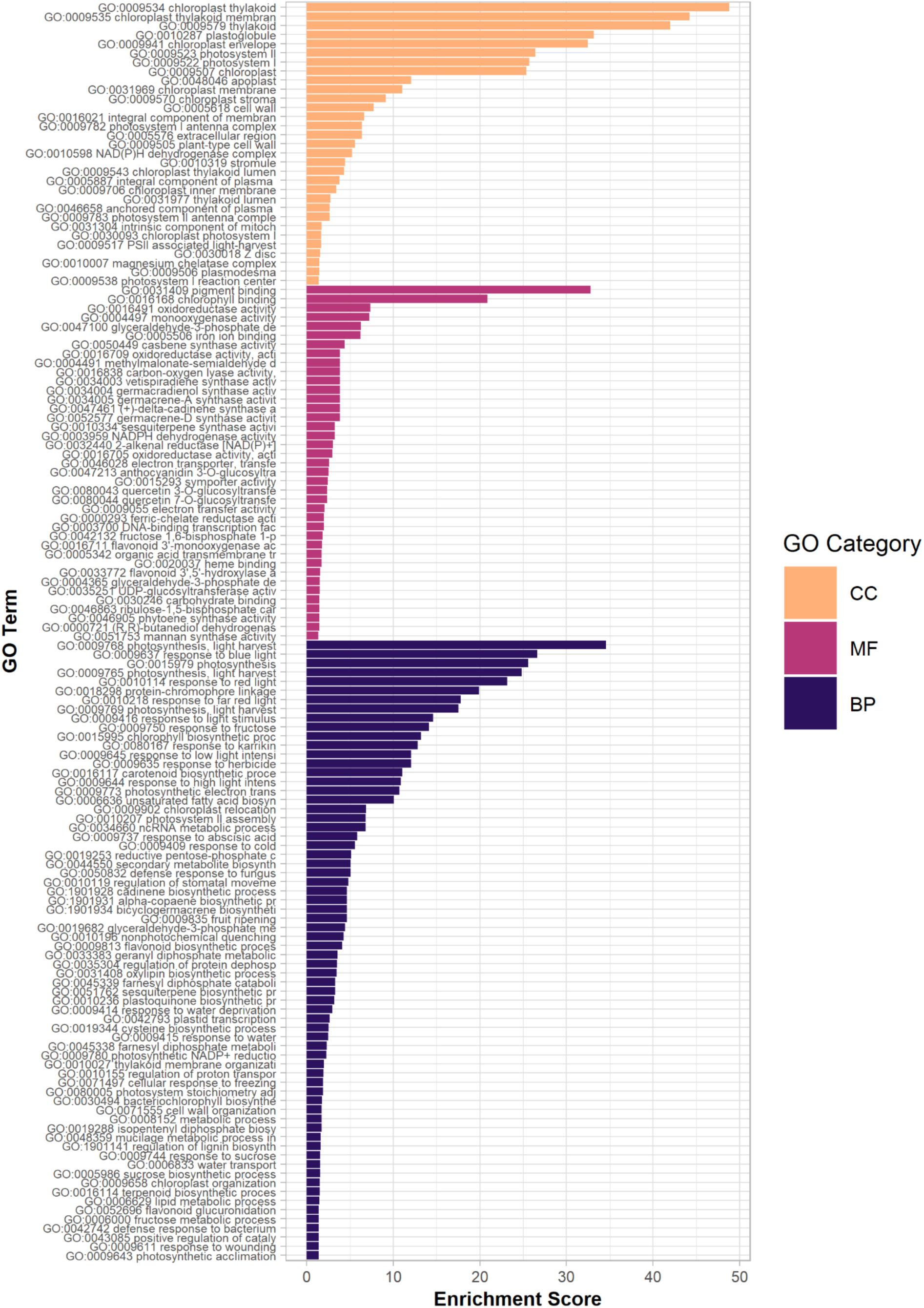
All enriched gene ontology terms for the differentially expressed genes shared in the responses to inoculation with *P. syringae* pv. *syringae* B728a (*Psy*) and *P. syringae* pv. *tabaci* ATCC11528 (*Pta)* at 5 hpi. GO terms were considered enriched for an adjusted *p* value<0.05. CC, cellular component; MF, molecular function; BP, biological processes.

**Supplemental Figure 4.**
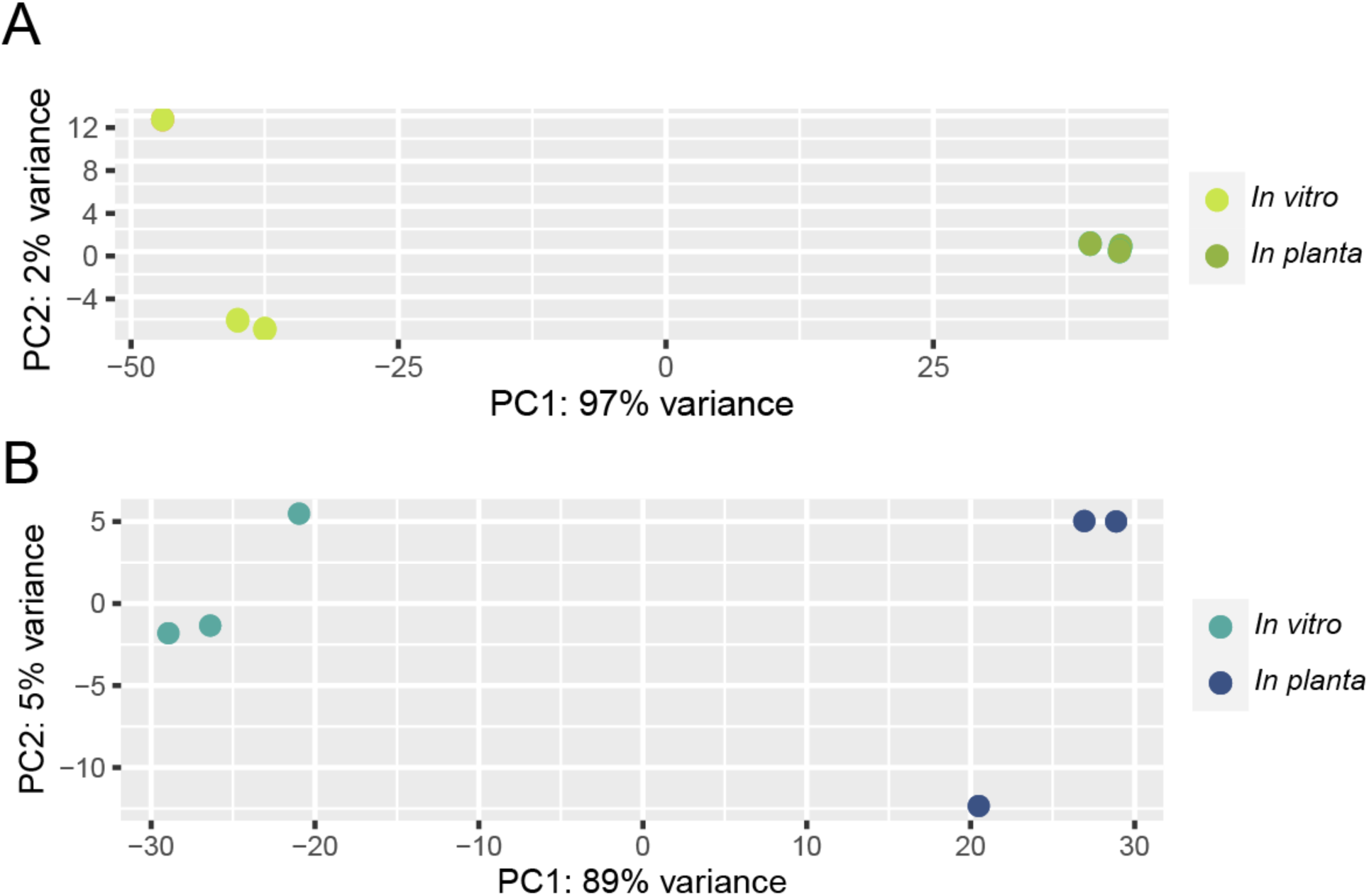
A principal component analysis plot depicting the three biological replicates for each of the four conditions sampled in the RNA-seq analysis of A) *P. syringae* pv. *tabaci* ATCC11528 (*Pta)* and B) *P. syringae* pv. *syringae* B728a (*Psy*).

**Supplemental Figure S5.**
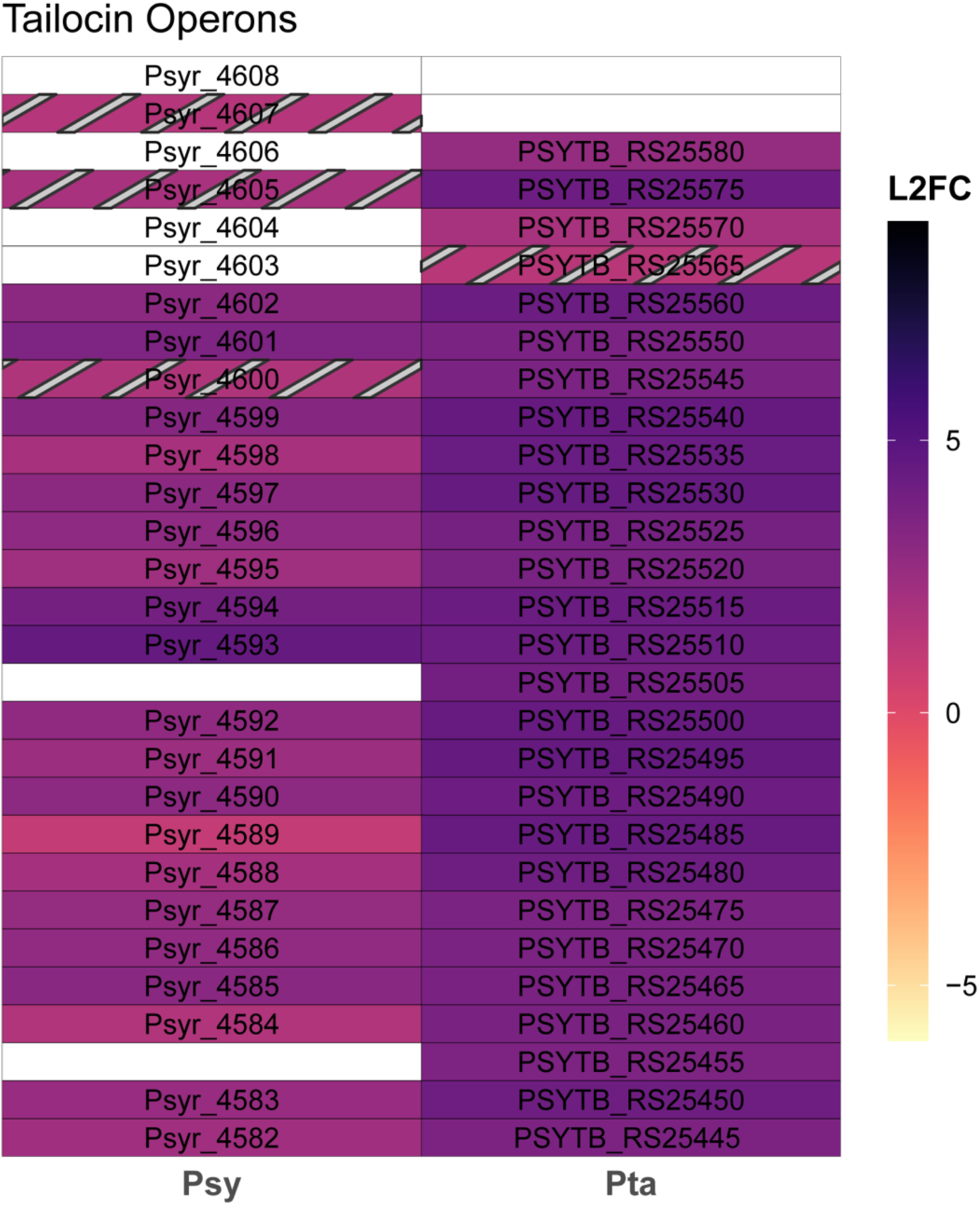
Differential expression of select tailocin loci in *P. syringae* pv. *syringae* B728a (*Psy*) and *P. syringae* pv. *tabaci* ATCC11528 (*Pta) in planta* based on an adjusted p value<0.01 and |log2 fold change|>2 . Striped boxes indicate genes that are significant for different thresholds: adjusted p value<0.05 and no minimum log2 fold change (L2FC). White boxes indicate no ortholog present or, when labeled with a locus ID, orthologs that are not differentially expressed for either set of thresholds.

**Supplemental Figure 6.**
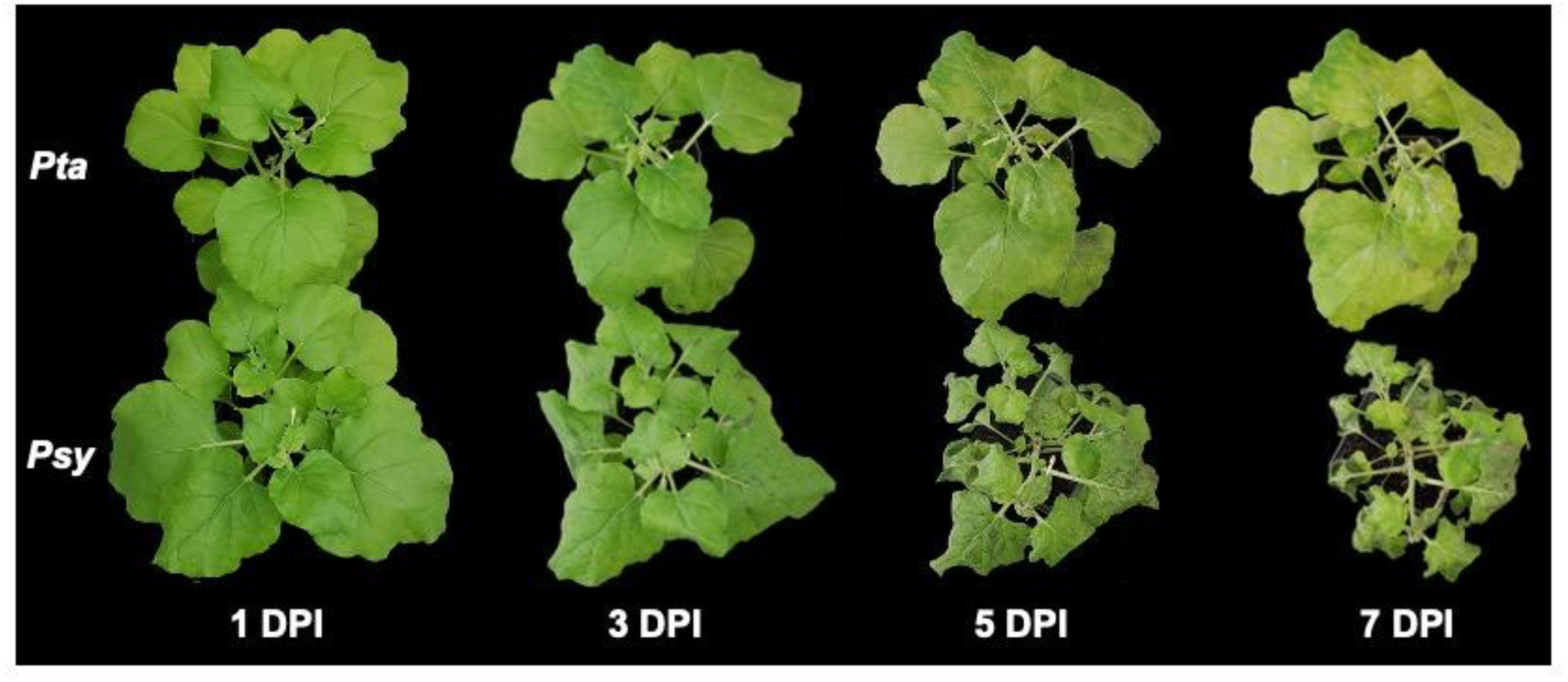
Infection time course comparing *P. syringae* pv. *syringae* B728a (*Psy*) and *P. syringae* pv. *tabaci* ATCC11528 (*Pta*) symptom development in *N. benthamiana*. Eight-week post seeding plants were held at 100% humidity for one day and then inverted submerged and gently agitated for 30 seconds in suspensions of *Pta* or *Psy* at 1×10^5 CFU/mL with 0.02% Silwet L77 . Plants were kept at 100% humidity under a 12 hour light/dark cycle and at ambient room temp and disease symptom progression was followed for 7 days.

**Supplemental Figure 7.**
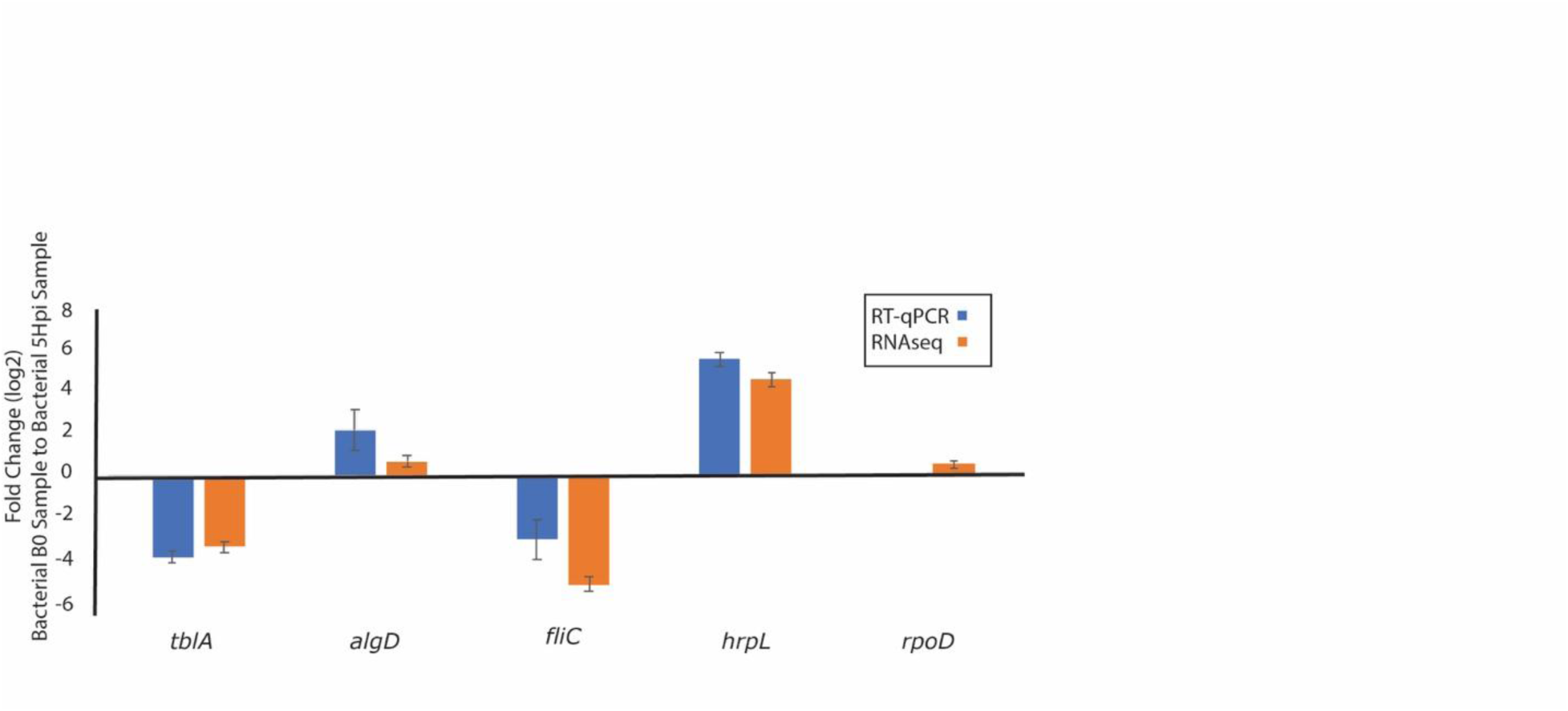
RT-qPCR Comparison of Virulence Gene Expression in *P. syringae* pv. *tabaci* ATCC11528. Samples were taken as per RNAseq comparisons, with B0 representing bacteria resuspended prior to infection and with sampling of leaf tissue at 5 hours past infection. Values are the result of three biological replicate experiments, and comparisons of Log2 fold change for each locus is shown from the RT-qPCR as well as from the RNAseq data. Error bars represent ± 1 standard error.

**Supplemental Figure S8:**
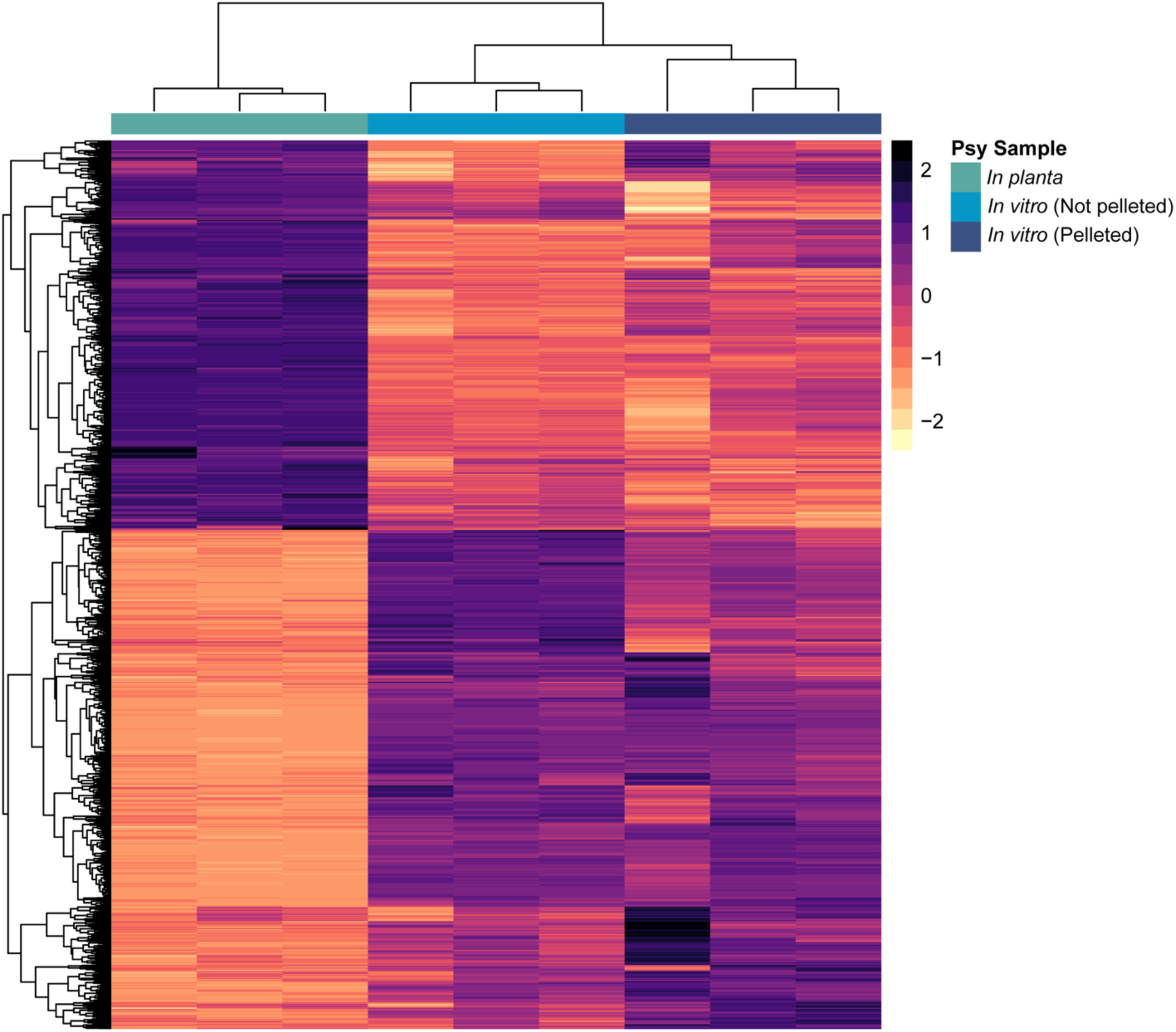
Effect of Centrifugation on *In Planta* Expression Calls for *P. syringae* pv. *syringae* B728a. RNAseq analyses were performed as described in Materials and Methods, but comparing gene expression patterns for three replicates of *Psy in planta* at T5 with patterns from three replicates of *Psy* at T0 that were either resuspended from agar (*in vitro* not pelleted) and those that were pelleted after resuspension (*in vitro* pelleted).

